# Conformational plasticity of a BiP-GRP94 chaperone complex

**DOI:** 10.1101/2024.02.01.578445

**Authors:** Joel Cyrille Brenner, Linda Zirden, Yasser Almeida-Hernandez, Farnusch Kaschani, Markus Kaiser, Elsa Sanchez-Garcia, Simon Poepsel, Doris Hellerschmied

## Abstract

Hsp70/Hsp90-chaperones and their regulatory co-chaperones are critical for maintaining protein homeostasis. GRP94, the sole Hsp90-chaperone in the secretory pathway of mammalian cells, is essential for the maturation of important secretory and transmembrane proteins. Without the requirement of co-chaperones, the Hsp70-protein BiP controls regulatory conformational changes of GRP94 – the structural basis of which has remained elusive. Here, we biochemically and structurally characterize the formation of a BiP-GRP94 chaperone complex and its transition to a conformation expected to support the loading of substrate proteins from BiP onto GRP94. BiP initially binds to the open GRP94 dimer via an interaction interface that is conserved among Hsp70/90 paralogs. Subsequently, binding of a second BiP protein stabilizes a semi-closed GRP94 dimer, thereby advancing the chaperone cycle. Our findings highlight a fundamental mechanism of direct Hsp70/90 cooperation, independent of co-chaperones.

## Introduction

Cellular protein homeostasis depends on the coordinated activity of molecular chaperones^1^. To this end, the Hsp70 and Hsp90 chaperone families are critical to maintain the integrity of the cellular proteome^2–4^. They are highly conserved across all domains of life, with eukaryotic cells having evolved compartment-specific Hsp70/90 paralogs. About 30% of the proteome in eukaryotic cells is processed in the secretory pathway, where the highly abundant BiP and GRP94 chaperones serve as the exclusive Hsp70/90 chaperone system^5–8^. BiP and GRP94 safeguard protein homeostasis in the endoplasmic reticulum (ER)^9–11^, in post-ER compartments^12,13^, and in the extracellular space^14–17^. BiP engages with exposed hydrophobic residues of newly synthesized or misfolded proteins and accordingly serves a broad substrate spectrum^18^. GRP94 is a more specialized chaperone, critical for the quality control of a subset of secretory and transmembrane proteins, including key growth factors and immune signaling proteins^19^. Previous work on Hsp70 and Hsp90 proteins from different organisms and different organelles established a role for Hsp70/BiP proteins in early substrate folding steps and assigned Hsp90/GRP94 proteins a downstream role in substrate maturation and activation^20–23^. A direct interaction between Hsp70 and Hsp90 proteins is critical for substrate protein handover^24–27^. Mutations in conserved Hsp70:Hsp90 interface residues lead to impaired substrate protein folding and growth defects in yeast^24,26,28,29^. To understand protein homeostasis in the secretory pathway, a detailed molecular understanding of the collaboration between GRP94 and BiP is critical.

GRP94, like other Hsp90 proteins, consists of an N-terminal ATPase domain (NTD), a middle domain (MD), and a C-terminal dimerization domain (CTD); all connected by flexible linkers^30^ (**Figure 1a**). Hsp90 proteins work as dimers, constitutively dimerized via their CTDs^2^. Additional dimerization of the NTDs is subject to regulation, where dynamic opening and closing via intermediate states drive substrate protein processing ^2^. Crystal structures of GRP94 show the chaperone in two extreme conformations – an open twisted V ^31^ and a fully closed conformation^32^ (**Figure 1a**). Unique features of GRP94 within the Hsp90 family include its N-terminal ER targeting sequence (1-21) and the pre-N domain (22-72), required for substrate protein maturation^32^ and important for the regulation of dimer closure and ATPase activity^31,32^ as well as an insertion in its ATP lid^33^. A truncated GRP94 protein, lacking the signal sequence and pre-N domain (GRP94 Δ72), shows increased ATPase activity compared to full-length GRP94, owing to accelerated ATP-driven dimer closure^32^. Cytosolic Hsp90 proteins rely on co-chaperones to promote the conformational cycle of opening and closing of the NTDs^2^. For GRP94 only two co-chaperones have been described, with a role in conferring substrate selectivity^34,35^. Notably, previous work has demonstrated that BiP acts as a closure-accelerating partner chaperone of GRP94^36,37^. BiP is an Hsp70-type chaperone with an N-terminal nucleotide-binding domain (NBD) and a C-terminal substrate binding domain (SBD), which is further subdivided into a beta-sheet rich base (SBDβ) and an alpha-helical lid (SBDα)^4,5^ (**Figure 1a**). An inter-domain linker between BiP_NBD_ and BiP_SBD_ provides the basis for allosteric regulation. ATP binding and hydrolysis of the BiP_NBD_ regulates the conformation of the BiP_SBD_, with the ATP-bound open and the ADP-bound closed conformation representing the most extreme states (**Figure 1a**)^38–41^. The SBD-open state is characterized by a high on/off rate for substrate proteins, and an interaction interface between BiP_NBD_ and BiP_SBD_^38–41^. It is therefore also referred to as the domain-docked conformation. Conversely, the domain-undocked conformation is stabilized by substrate protein bound to the closed BiP_SBD_, which is entirely detached from the BiP_NBD_^39^. In its collaboration with GRP94, BiP serves two major roles that are closely connected: BiP delivers substrates to GRP94 and accelerates GRP94 dimer closure^36,37,42^. Together, these activities should facilitate substrate loading onto GRP94. A recent high-resolution cryo-electron microscopy (cryo-EM) structure provided important insights into substrate transfer between the cytosolic Hsp70/90 chaperone systems. In the so-called loading complex, resulting from interactions between Hsp70, Hsp90, the co-chaperone Hop, and a substrate protein, Hsp90 adopts a semi-closed conformation, where the NTDs have dimerized, but are not yet in an ATPase-competent state^24^. Importantly, Hop organizes the loading complex by bridging Hsp70 and Hsp90, stabilizing the semi-closed Hsp90 conformation, and extending the substrate interaction interface of the complex^24^.

**Figure 1.**
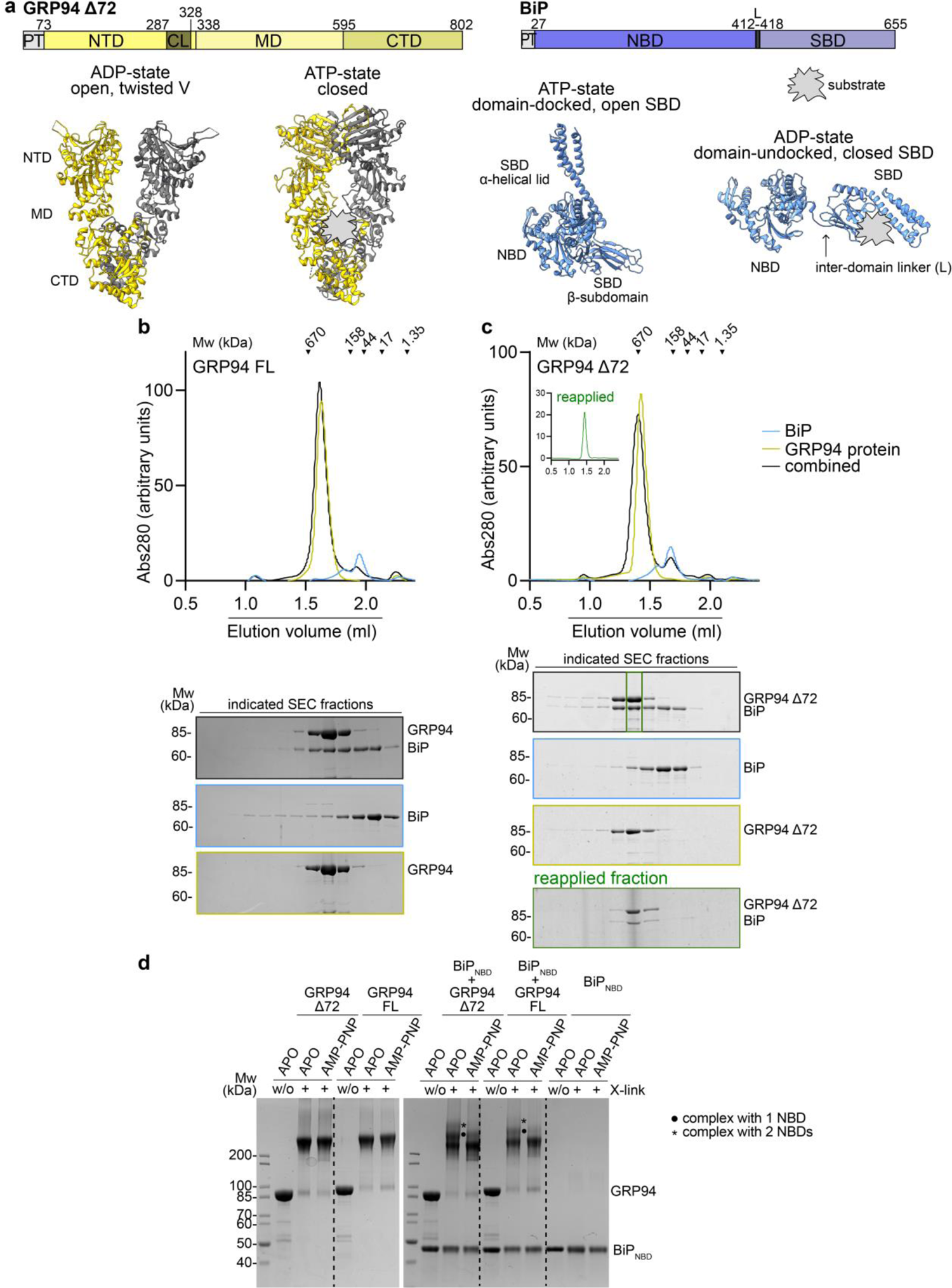
Biochemical characterization of a BiP-GRP94 complex. (**a**) Crystal structures of the open GRP94 dimer (PDB ID: 2O1V), the closed GRP94 dimer (PDB ID: 5ULS), domain-docked BiP (PDB ID: 5E84), and a model of BiP based on the domain-undocked conformation of its homolog DnaK (PDB ID: 2KHO). NTD: N-terminal domain, MD: middle domain, CTD: C-terminal domain, NBD: nucleotide-binding domain, SBD: substrate-binding domain. (**b**) Analytical SEC and SDS-PAGE analysis of complex formation between GRP94 full-length (FL) and BiP. (**c**) Same analysis as in (b) for GRP94 Δ72 and BiP. The fraction containing a BiP-GRP94 Δ72 complex (green box) was reapplied to the SEC column. (**d**) SDS-PAGE of glutaraldehyde crosslinking reactions containing GRP94 Δ72 or FL protein and BiP_NBD_ in the presence and absence of AMP-PNP.

Despite the high conservation and functional similarities between Hsp70/90 systems, across species and organelles, their regulation clearly differs^2,19,43^. Most prominently, bacteria as well as organellar compartments lack a bridging co-chaperone like Hop. How BiP is structurally integrated into the GRP94 conformational cycle, in the absence of bridging co-chaperones, has remained elusive. Here we describe two distinct states of the BiP-GRP94 complex. In the ‘pre-loading complex’, a direct interaction between BiP and the open GRP94 dimer is established via a highly conserved docking site. GRP94 and BiP subsequently move on to a loading complex conformation previously reported for the cytosolic Hsp70/90/Hop system ^24^ even in the absence of any co-chaperone organizing their interaction. Our biochemical and structural data provide insights into a conserved mechanism of chaperone-complex formation and priming for substrate transfer from Hsp70/BiP to Hsp90/GRP94 machineries.

## Results

### A stable complex between BiP and GRP94

To biochemically and structurally characterize the formation of a BiP-GRP94 complex, we reconstituted the interaction between the two chaperones *in vitro*. We recombinantly expressed and purified Strep-tagged GRP94 (residues: 23-802) and His_6_-tagged BiP (residues: 27-655), both lacking their N-terminal ER-targeting signal sequences. In agreement with previous studies, BiP displayed a mixture of oligomeric and monomeric species in size exclusion chromatography (SEC) (**Figure 1b**)^37,44^. Nevertheless, we could perform analytical SEC studies to monitor complex formation with GRP94. Our data show that BiP elutes in higher molecular weight fractions together with the GRP94 full-length (FL) wild-type protein (**Figure 1b**). We did not include nucleotides in our interaction studies of full-length BiP and GRP94, since GRP94 exclusively binds the domain-undocked, ADP-bound conformation of BiP^37^. To study the interaction between BiP and different states of the GRP94 dimer (controlled by nucleotide binding and hydrolysis), we turned to GRP94 ATP binding and hydrolysis mutants. The GRP94 wild-type and D149N nucleotide binding mutant protein are expected to preferentially populate an open conformation, while the E103A ATP-hydrolysis mutant is expected to reside mainly in a closed state^32,45^. Compared to GRP94 wild-type protein we observed a minor increase in BiP co-eluting with GRP94 D149N (**Extended Data Figure 1a**), while GRP94 E103A showed slightly reduced interaction with BiP (**Extended Data** Figure 1a). These results indicated that the closed GRP94 state is less favorable for BiP binding.

We also tested the interaction of BiP with a truncated form of GRP94, which lacks the pre-N domain (referred to as GRP94 Δ72) (**Figure 1a**). GRP94 Δ72 has previously been shown to functionally interact with BiP, specifically with its NBD^36,37^. In analytical SEC experiments, we observed a stable complex between BiP and GRP94 Δ72, illustrated by the fact that upon reapplying the complex to the SEC column, BiP and GRP94 Δ72 still co-eluted in a single peak (**Figure 1c**). To further assess the effect of the GRP94 nucleotide-bound state on complex formation we used a BiP_NBD_ construct, which irrespective of its nucleotide state, retains GRP94 binding^36,37^ (**Figure 1a**). In analytical SEC experiments, BiP_NBD_ did not co-elute with GRP94 Δ72 (**Extended Data** Figure 1b). However, in chemical crosslinking experiments using glutaraldehyde, we could stabilize a complex between the BiP_NBD_ and GRP94 FL or GRP94 Δ72. In the presence of excess amounts of AMP-PNP, which shifts the GRP94 dimer to the closed conformation ^32^, the interaction is strongly reduced (**Figure 1d** and **Extended Data** Figure 1c). The nucleotide dependence of the interaction is recapitulated for the truncated GRP94 Δ72 construct, with an even more pronounced band corresponding to a GRP94 Δ72 dimer with two BiP_NBD_ molecules bound (**Figure 1d** and **Extended Data** Figure 1c). Taken together, we established a protocol to isolate a stable BiP-GRP94 complex, favoring the truncated form of GRP94 in the absence of ATP(-analogues).

### Hydrophobic tagging to establish chaperone-substrate complexes in vitro

To better understand how GRP94 as well as a BiP-GRP94 complex interact with substrate proteins, we established interaction studies with a model substrate. BiP and GRP94 have previously been shown to interact with a misfolded HaloTag2 (HT2)-based model substrate in HEK293 cells^46,47^. The HT2 domain is a self-labeling protein tag, which covalently conjugates to chloroalkanes^48^. In a process called hydrophobic tagging, HT2 covalently conjugates to a hydrophobic tag, which induces HT2 misfolding in cells (**Figure 2a**)^49–52^. We characterized the misfolding process *in vitro* by limited proteolysis using trypsin. Upon incubation of HT2 with HyT36 the protein is more susceptible to tryptic digest, indicating a misfolded state (**Figure 2b**). Notably, HT2 misfolding is dependent on the covalent conjugation of HyT36 to HT2. In the presence of a control compound, HyT36(-Cl), which cannot conjugate to HT2, the protein remains stable; excluding a non-specific destabilizing effect (**Figure 2b**). In circular dichroism (CD) spectroscopy experiments we analyzed the kinetics of misfolding (**Figure 2c**). The alpha-helical content of HT2 (measured by an increase in signal at 222 nm) reaches its minimum at ∼30 min incubation at 37 °C, revealing experimental conditions that can be applied in chaperone-substrate interaction studies. At 25 °C, HT2 conjugated to HyT36 remains stable (**Figure 2c**), thereby showing the temperature-dependence of hydrophobic tagging-induced misfolding. Conjugation of HT2 to the control compound 6-chlorohexanol, a HaloTag ligand without any functional group, did not affect the stability of the HT2 protein at 25 °C or 37 °C (**Figure 2c**).

**Figure 2.**
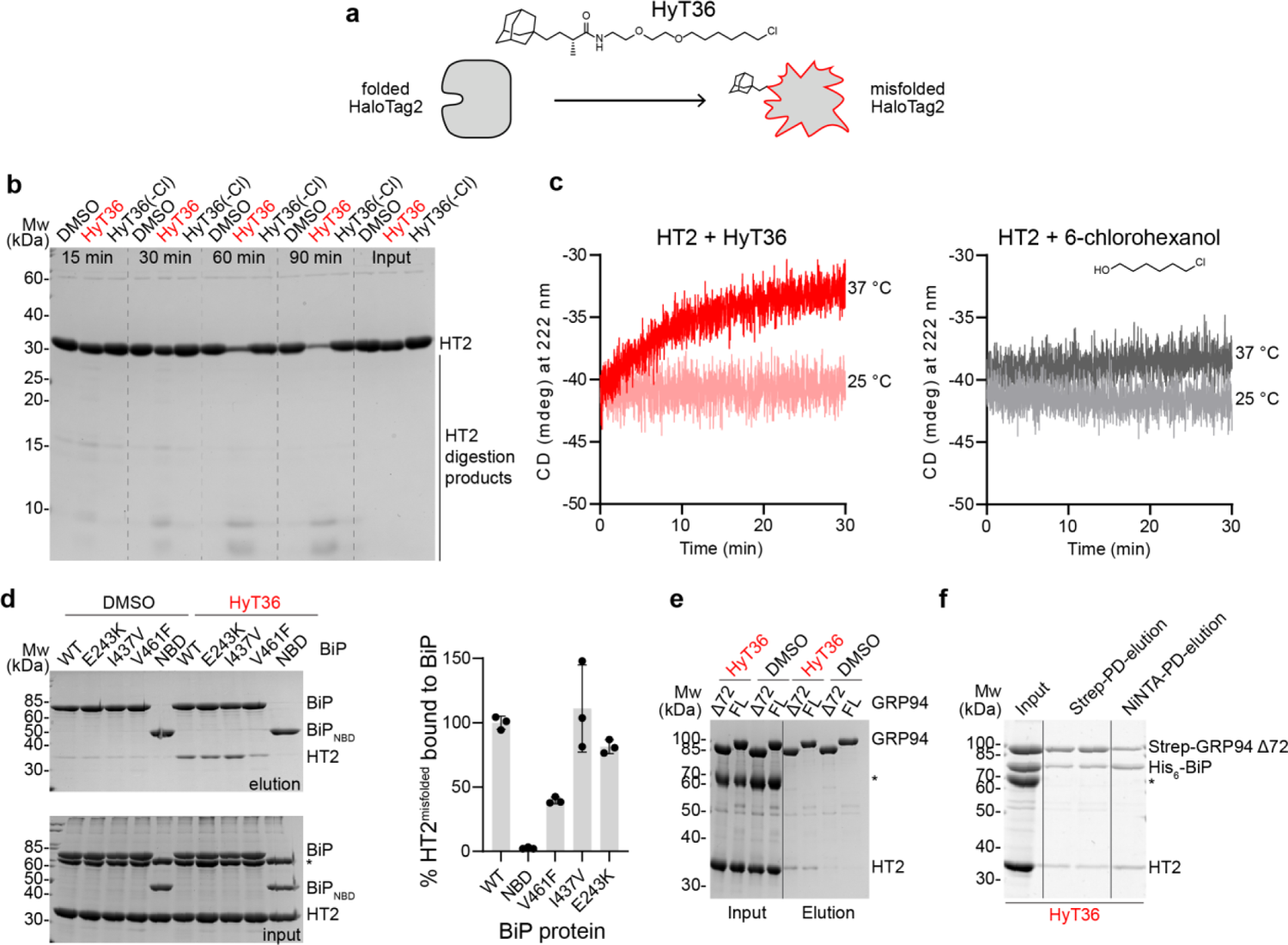
Hydrophobic tagging to establish a BiP-GRP94-substrate complex. (**a**) Cartoon model of HaloTag2 (HT2) misfolding upon covalent conjugation to HyT36. (**b**) Limited proteolysis with trypsin to assess the folding state of HT2 conjugated with HyT36, incubated with the HyT36(-Cl) control, or DMSO. (**c**) CD spectroscopy of HT2 conjugated to HyT36 or 6-chlorohexanol. Kinetic measurements at the indicated temperatures are shown. (**d**) Pull-down experiments to study the interaction of His_6_-BiP with misfolded HT2-HyT36 compared to DMSO control. For quantification data were normalized to the WT BiP elution sample. (**e**) Same analysis as in (d) for GRP94 FL and GRP94 Δ72. (**e**) Sequential pull-down of Strep-GRP94 Δ72 and His_6_-BiP to isolate a GRP94 Δ72-BiP-HT2^misfolded^ complex. The asterisk indicates BSA in the interaction buffer.

The interaction of Hsp70-type chaperones with misfolded substrate proteins is well characterized^53–55^. For *in vitro* experiments, substrate proteins are often destabilized with non-specific chemical denaturing agents. Upon dilution of the denaturing agent, chaperones can then be added to study chaperone-substrate interactions^56,57^. With the *in vitro* hydrophobic tagging system, protein misfolding can however be induced directly in the presence of the Hsp70-protein BiP. This setup provides BiP direct access to misfolded HT2 states as they arise. We conjugated HyT36 to HT2 by simple mixing of the two components, followed by the addition of BiP and ATP, and finally incubation at 37 °C to induce HT2 misfolding. Under these conditions, misfolded HT2 (HT2^misfolded^) is directly bound by BiP, as observed in pull-down assays (compare DMSO to HyT36 condition, **Figure 2d**). Deletion of the BiP substrate binding domain entirely abolished BiP-HT2^misfolded^ interaction (**Figure 2d** – BiP_NBD_). A previously described substrate-binding mutant, BiP V461F^44,58,59^ showed reduced interaction with HT2^misfolded^. We also tested BiP I437V, a mutation described to stabilize the closed (domain-undocked) conformation^39^ and BiP E243K, a mutation at the BiP_NBD_:BiP_SBD_ interface, described to have reduced interaction with GRP94^37^. While BiP I437V did not consistently affect the binding of the substrate, BiP E243K showed a trend towards slightly reduced substrate binding (**Figure 2d**). We then performed similar experiments with full-length GRP94 and GRP94 Δ72, which also bound to HT2 upon misfolding (**Figure 2e**). Finally, we performed a sequential pull-down experiment to assess whether a HT2^misfolded^/substrate-loaded complex containing BiP and GRP94 Δ72 can be formed. We first pulled on Strep-tagged GRP94 Δ72, followed by His_6_-tagged BiP to isolate complexes that contained both chaperones. In the final elution fraction, we detected HT2, demonstrating the formation of a GRP94 Δ72-BiP-HT2^misfolded^ complex (**Figure 2f**).

### Assessing the topology of a BiP-GRP94 complex by crosslinking mass spectrometry (XL-MS)

In order to gain initial structural insight into a BiP-GRP94-substrate complex we performed XL-MS experiments. We reconstituted complexes between BiP and GRP94 Δ72 in the presence and absence of HT2^misfolded^. Upon crosslinking with DSBU, SDS-PAGE gel bands containing high molecular weight protein complexes were analyzed by MS (**Extended Data Table 1, Figure 3a**, and **Extended Data Figure 2a**). The overall crosslinking pattern of BiP and GRP94 Δ72 did not change with the presence of HT2, however, we could also not detect any crosslinks to the misfolded HT2 substrate protein (**Figure 3a** and **Extended Data Figure 2a**).

**Figure 3.**
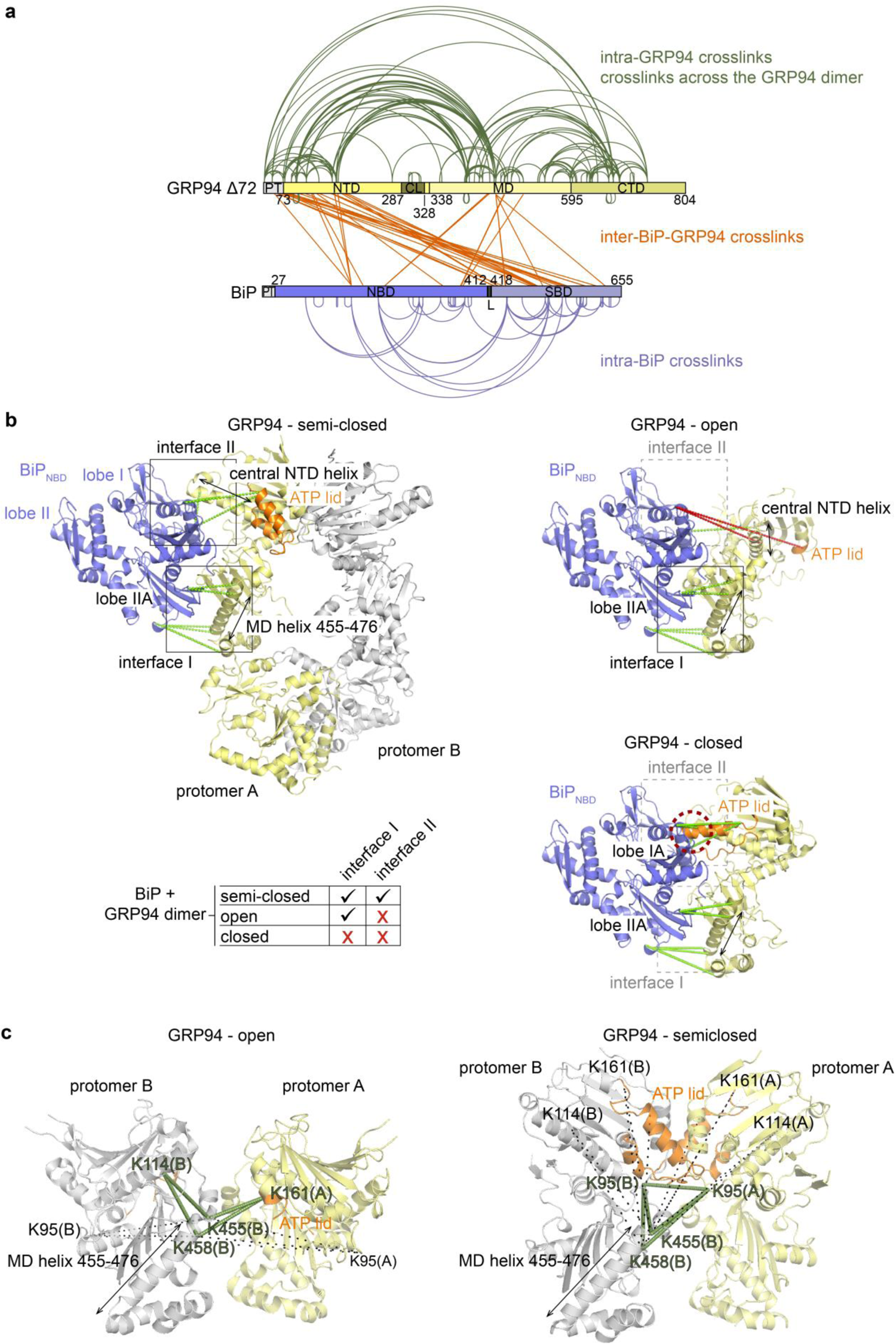
Crosslinking mass spectrometry outlines the topology of a BiP-GRP94 Δ72 complex. (**a**) Map of crosslinks identified in the BiP-GRP94 Δ72-HT2 sample. (**b**) *Left panel:* BiP_NBD_-GRP94 crosslinks mapped onto a homology model of BiP-GRP94 on the basis of the cytosolic loading complex (PDB ID: 7KW7). *Right panel:* Same crosslinks mapped onto GRP94 proteins in the open (2O1V-chain A) and closed (5ULS-chain A) conformation aligned to the semi-closed conformation model shown in the left panel. Structures are aligned based on the MD and CTD of GRP94 protomer A. (**c**) Intra-GRP94 crosslinks that are selectively supported in the open (PDB ID: 2O1V) or semi-closed (model also shown in b) GRP94 dimer. For simplicity, only MD and NTD are shown. Crosslinks shown as black dashed lines are > 30 Å. NTD: N-terminal domain, MD: middle domain, CTD: C-terminal domain, NBD: nucleotide-binding domain, SBD: substrate binding domain, PT: purification tag, CL: charged linker, L: linker.

For the following analysis, we focused on the sample containing all three proteins. The observed crosslinks between BiP and GRP94 Δ72 suggest that the two chaperones adopt a similar conformation as their cytosolic paralogs, Hsp70 and Hsp90, in the previously described loading complex^24^. Figure 3b shows a homology model of BiP and GRP94 on the basis of the cytosolic loading complex (PDB ID: 7WK7). All detected BiP_NBD_-GRP94 crosslinks map to two interaction interfaces (I and II) (**Figure 3b, Extended Data Table 1**), matching interactions described for the cytosolic loading complex. The crosslinks outlining interface I link the central GRP94_MD_ helix to BiP_NBD_ lobe IIA. The crosslinks that indicate the formation of interface II link the central GRP94_NTD_ helix and the ATP lid to the BiP_NBD_ lobe I. Notably, interface II crosslinks are only compatible with the presence of a semi-closed conformation of GRP94, as reported for Hsp90 in the cytosolic loading complex^24^. The open GRP94 conformation exclusively allows for interactions within interface I, while interface II residues are too far apart (**Figure 3b**). In the GRP94 closed conformation the ATP lid would clash with residues of BiP_NBD_ lobe IA, preventing an efficient interaction. This effect is consistent with our biochemical data showing that the AMP-PNP-bound and thus closed GRP94 dimer does not interact with the BiP_NBD_ (**Figure 1d**).

To define the conformation of GRP94, we therefore next analyzed intra-protein crosslinks and crosslinks across the GRP94 dimer. Within the GRP94_CTD_ and GRP94_MD_, 46/50 unique crosslinks agree with published structures, validating the quality of the crosslinking data (**Extended Data Figure 2b**); however, these crosslinks cannot distinguish between different GRP94 conformations. We therefore analyzed crosslinks between GRP94_NTD_s and GRP94_MD_s in more detail. On the one hand, we found crosslinks from the GRP94_NTD_ ATP lid (residues K161 and K168) and the central NTD helix (residue K114) to GRP94_MD_ helix 455-467. These are exclusively compatible with an open twisted V conformation (**Figure 3b**). Upon rotation of the GRP94_NTD_s to the semi-closed conformation, the central NTD helix and the ATP lid move away from the GRP94_MD_. At the same time, repositioning of the K95 residues now supports a crosslink between the two across the GRP94 dimer, as identified in our sample (**Figure 3b**). Collectively, the data suggest that our sample is heterogeneous and may contain open (twisted V) and semi-closed GRP94 dimers in complex with BiP.

Crosslinks found within the BiP_SBD_ confirm that when in complex with GRP94, it resides in the closed conformation (**Extended Data Figure 2c**). The closed BiP_SBD_ is fully dissociated from the BiP_NBD_ and free to move around the inter-domain linker^39,60^. The 19 identified BiP_SBD_-GRP94 crosslinks suggest two positions for the BiP_SBD_ relative to GRP94. Two crosslinks (GRP94_MD_ residue 458 to BiP_SBD_ residues 446/523) are compatible with the conformation of the Hsp70 SBDβ resolved in the cytosolic loading complex^24^ (**Extended Data Table 1**). The majority of GRP94-BiP_SBD_ crosslinks however place the BiP_SBD_ in the proximity of the GRP94_NTD_ and suggest an interaction with the N-terminal GRP94 Δ72 purification tag (**Extended Data Table 1** and **Figure 3a, Extended Data** Figure 2a).

### BiP binds to the open GRP94 dimer – formation of the pre-loading complex

To study the topology and conformations of a BiP-GRP94 Δ72 complex, we performed single-particle negative-stain electron microscopy (EM). Even though we could not detect any crosslinks to HT2, we hypothesized that the misfolded substrate may stabilize or support the formation of the BiP-GRP94 chaperone complex and we therefore included HT2^misfolded^ in our sample preparation. To establish a BiP-GRP94 Δ72-HT2^misfolded^ complex, we first incubated BiP with HT2 in the presence of HyT36 and ATP. We then added GRP94 Δ72 to bind the preassembled BiP-HT2^misfolded^ complex. To enrich the multi-chaperone-substrate complex, we performed Gradient Fixation (GraFix) ^61^, which resulted in the separation of three major complex species (**Extended Data Figure 3a**). Based on their sizes, we identify a GRP94 dimer (complex 0) and two complexes compsed of a GRP94 dimer, HT2 and one or two BiP molecules, respectively (complexes I and II). GraFix fraction 12 contained the largest amounts of complex I and II with little amount of free GRP94 dimer and was therefore used for negative-stain EM (**Extended Data Figure 3a**). As expected, reference-free 2D classifications and *ab initio* 3D reconstructions indicated sample heterogeneity (**Extended Data Figure 3b** and 4). Nevertheless, the *ab initio* 3D reconstructions in comparison to known GRP94 structures showed two predominant complexes corresponding to a GRP94 dimer with one (complex I) and two (complex II) BiP proteins bound (**Extended Data** Figure 4).

In the complex I reconstruction, we identified a density corresponding to a GRP94 dimer with a gap between the NTDs (**Figure 4a**). The additional density attached to GRP94 protomer A resembles a BiP_NBD_ domain with a characteristic globular shape outlining lobes I and II, and an indentation corresponding to the nucleotide-binding cleft (**Figure 4a**). Accordingly, we chose an open twisted V GRP94 Δ72 crystal structure reported by Dollins et al. (PDB ID: 2O1V ^31^) and the structure of the isolated BiP_NBD_ (PDB ID: 6DWS) as the starting structures for molecular modeling. After an initial rigid body fit to place the structures into the density, we performed a map-fitting refinement of the complex with Rosetta ^62,63^. Both the *total score* and the *elect_dens_fast* score values of the obtained models, negatively correlate with the real-space correlation coefficient (RSCC) values (**Extended Data Figure 5a**), probably because the models could fit most of the density (**Figure 4b**). We presume that the BiP_SBD_ is rather flexible in this complex and therefore largely not visible in the EM map. A density close to the GRP94_NTD_ protomer B, not accounted for in our model, may correspond to the BiP inter-domain linker and/or parts of the BiP_SBD_ (**Figure 4ab**). We refer to this state of the BiP-GRP94 complex as the pre-loading complex, a conformation that exists prior to the dimerization of the GRP94_NTD_s i.e. before the full engagement of GRP94 with substrate protein. As expected, for a BiP-GRP94 complex with GRP94 in the open twisted V conformation, the pre-loading complex model is supported by our crosslinking data outlining interface I, while interface II has not formed yet (**Extended Data Figure 5b** and **Figure 3b** for comparison). The pre-loading complex depicts the initial engagement of BiP with the open GRP94 dimer, the substrate-loading competent form.

**Figure 4.**
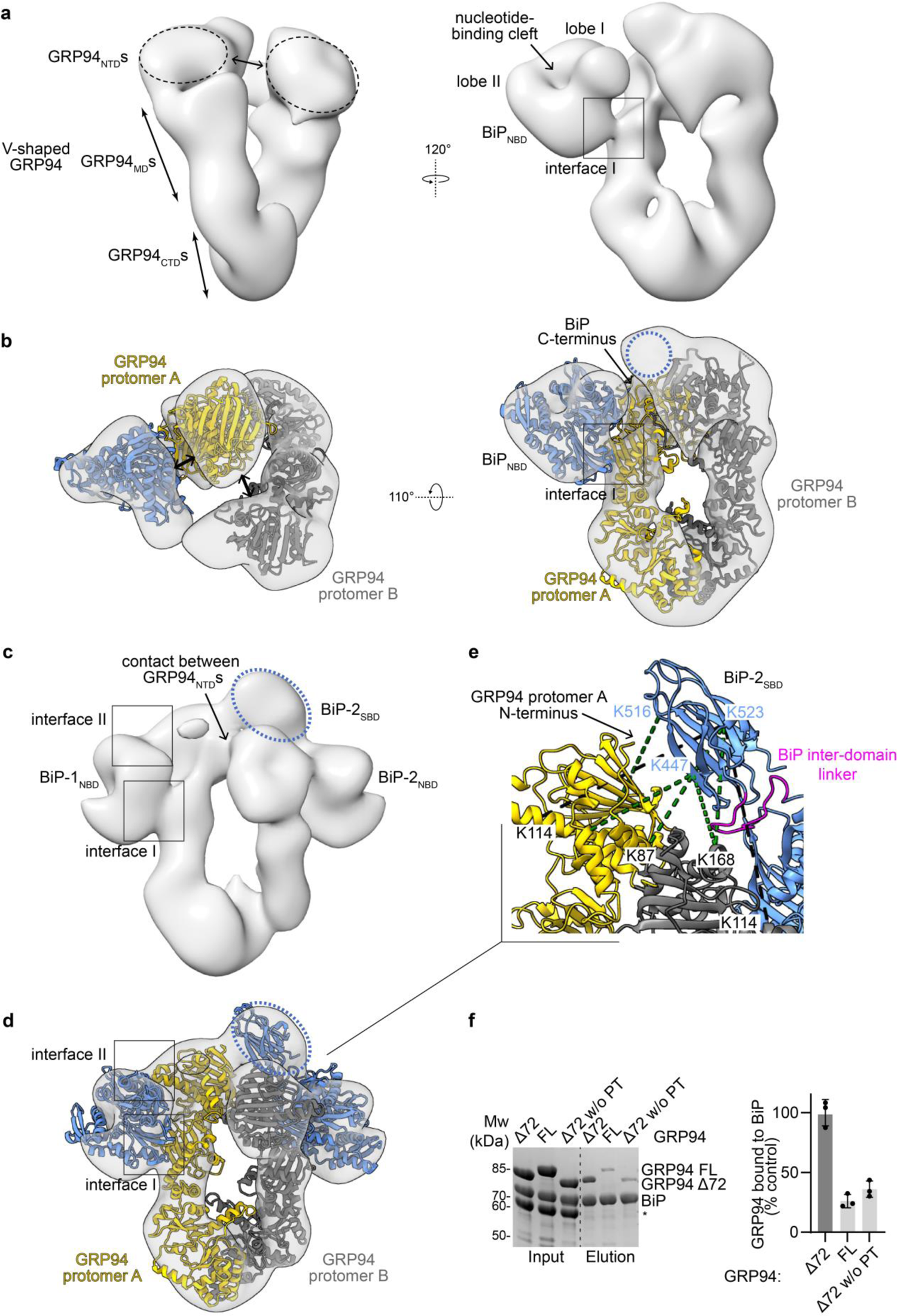
Structure of the BiP-GRP94 complex in two different conformations. **(a)** Negative-stain EM reconstruction and (**b**) molecular model of the BiP-GRP94 pre-loading complex. (**c**) Negative-stain EM reconstruction and (**d**) molecular model of the BiP-GRP94 loading complex. The dotted blue circles indicate the potential position of the BiP_SBD_. (**e**) Crosslinks identified between BiP_SBD_ and GRP94 Δ72 mapped onto the loading complex structure. Green lines indicate crosslinks <35 Å, black lines >35 Å. The arrow indicates the N-terminus of GRP94 i.e. the attachment point of the purification tag. (**f**) *Left:* Ni-NTA pull-down analysis of His_6_-BiP with Strep-tagged GRP94 Δ72, Strep-tagged GRP94-FL, and a GRP94 Δ72 construct lacking the purification tag (w/o PT). See Extended Data Figure 6a for exact sequences. The asterisk indicates BSA in the interaction buffer. *Right:* Quantification of the pull-down experiment (n = 3, mean +/- sd).

### BiP bound to a semi-closed state of GRP94 – the loading complex conformation

In addition to the pre-loading complex, we obtained a reconstruction that indicated the binding of two BiP molecules to a GRP94 dimer (complex II) and largely resembled the cytosolic loading complex reported by Wang *et al.*^24^. Our *ab initio* reconstruction shows that the GRP94_NTD_s have established a contact at the center of the dimer, indicating a semi-closed conformation (**Figure 4c**). The NBDs of the two outlined BiP proteins furthermore established interactions with the GRP94_MD_s (interface I) and the GRP94_NTD_s (interface II). To initiate molecular modeling, we obtained a homology model of GRP94 in the semi-closed state using the loading complex structure of Hsp90 as a template (PDB ID: 7KW7^24^). For the BiP-1_NBD_ we used the same structure as for the pre-loading complex (PDB ID: 6DWS). Since we observed an additional density connected to the BiP-2_NBD_ in proximity to the GRP94_NTD_ of protomer A (**Figure 4c**), we included the BiP_SBD_ β-subdomain in our model. We used a homology model of an extended, domain-undocked conformation of BiP_NBD-SBD_ based on the structure of its *Escherichia coli* homolog DnaK (PDB ID: 2KHO)^64^. The individual models were placed into the EM reconstruction and structural refinement allowed us to fit the extended conformation of BiP-2_NBD-SBD_ into the density (**Figure 4d**). We refer to this state of the BiP-GRP94 complex as the loading complex, given the similarity to the previously reported Hsp70/90 structure (**Extended Data Figure 5c**)^24^. The fitting scores and the RSCC for the loading complex as compared to the pre-loading complex lack a clear correlation (**Extended Data Figure 5a**). This could be explained by the larger size and complexity of the loading complex model and by the fact that at a lower contour level, the EM reconstruction showed a density connected to BiP-1_NBD_ that is not accommodated by the final model (**Extended Data Figure 5a**). We did not observe any additional densities that could be assigned to the HT2^misfolded^ in either of the two EM reconstructions. This may be due to its flexibility, size, and resolution limits of negative-stain EM.

Notably, in both EM reconstructions, the BiP_SBD_ appears to extend towards the NTD of the opposing GRP94 protomer (**Figure 4a-d**). In the pre-loading complex, residual density in combination with the BiP_NBD_ C-terminus indicates the position of the BiP_SBD_ (**Figure 4b**). In the loading complex, the defined BiP-2_SBD_ extends towards the GRP94_NTD_ protomer A and an additional interaction between the BiP inter-domain linker and the GRP94_NTD_ of protomer B is formed (**Figure 4d, e**). This position of the BiP-2_SBD_ with respect to GRP94 is in agreement with our crosslinking data (**Figure 4e**). Crosslinks connecting the BiP_SBD_ to the, presumably flexible, N-terminal GRP94 purification tag would likely also agree with this conformation (**Figure 4e**). Upon close inspection of the purification tag sequence, we identified a hydrophobic stretch and hypothesized that it may serve as a BiP substrate mimetic (**Extended Data Figure 6a**). To address the contribution of the GRP94 purification tag to the interaction, we removed the tag by TEV protease cleavage and performed comparative pull-down analyses (**Figure 4f**). Upon removal of the GRP94 purification tag, we observed a reduction in GRP94 protein bound to BiP by ∼65%, suggesting that the tag contributes to the stability of the chaperone complex. In line with this, we also observed ∼75% less GRP94 full-length (FL) protein pulled down with BiP compared to GRP94 Δ72 (**Figure 4f**). The full-length protein, while carrying a Strep-tag, does not contain the predicted BiP substrate sequence (**Extended Data Figure 6a**).

The binding of the BiP_SBD_ to the GRP94 Δ72 purification tag contributes an additional interaction interface and we presume that it prohibits transition to the fully closed GRP94 conformation, which is expected to induce complex disassembly (**Figure 1d**). Taken together, the stabilization of the BiP-GRP94 Δ72 complex by this inadvertently engineered interaction allowed us to capture two chaperone complex transition states.

### BiP_NBD_-GRP94 interactions in the pre-loading and loading complex conformations

In the two obtained EM reconstructions, the preferred position of the BiP_SBD_ is determined by the GRP94 Δ72 purification tag. Given that in the domain-undocked state, the BiP_NBD_ acts independently of the BiP_SBD_, we further characterized BiP_NBD_-GRP94 interactions (**Figure 5a**). In the pre-loading complex, the BiP_NBD_ and GRP94 showed one major contact point with an interface area (GRP94_MD_:BiP_NBD_) of 1072.5 Å^2^ (**Figure 5b**). According to our model, the long GRP94_MD_ helix (residues 455-476) packs against the BiP_NBD_ lobe IIA, thereby forming electrostatic interactions involving BiP residues D238, N239, and E243, and GRP94 residues K462, K463, R466, and K467 (**Figure 5b**). Additionally, the tip of a GRP94_MD_ beta sheet inserts between BiP_NBD_ lobe IA and IIA. This interaction matches interface I in the previously described cytosolic loading complex^24^ (**Figure 5b**). Mutations at interface I impair the interaction and functional cooperation between Hsp70 and Hsp90 chaperones^25,26,28,37^. Consistently, BiP E243K (E218 in cytosolic Hsp70) and GRP94 K463E or K467E (K414 and K419 in cytosolic Hsp90) strongly reduce BiP-GRP94 Δ72 complex formation in analytical SEC studies and pull-down experiments (**Figure 5cd**). For all three mutant proteins, a ∼85% reduction in the amount of GRP94 Δ72 co-purified with BiP was observed in pull-down experiments (**Figure 5c**), highlighting the importance of this conserved interface.

**Figure 5.**
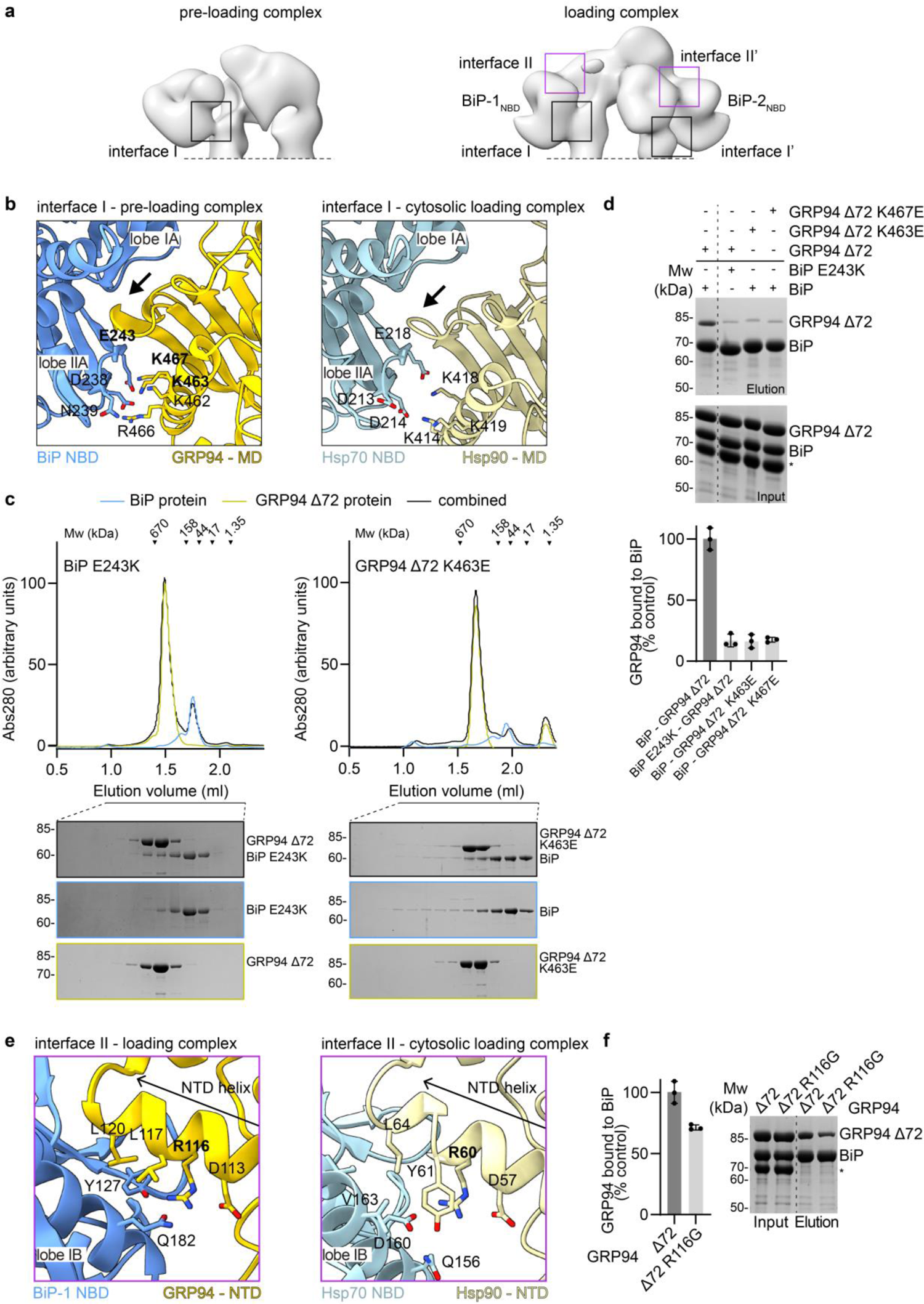
Interactions between BiP_NBD_ and GRP94. (**a**) Overview of interface I and II in the pre-loading and loading complex conformations. (**b**) Zoom-in into interface I of the BiP-GRP94 pre-loading complex. Interface I of the cytosolic (Hsp70/90) loading complex is shown for comparison. The arrow points to the GRP94_MD_ beta sheet loop inserted between BiP_NBD_ lobe IA and IIA. Residues in bold are used for mutational analysis. (**c**) Analytical SEC and SDS-PAGE analysis of complex formation between BiP and GRP94 Δ72 interface I mutants. (**d**) Ni-NTA pull-down analysis of His_6_-tagged BiP and Strep-tagged GRP94 Δ72 interface I mutants and quantification of the pull-down experiment (n = 3, mean +/- sd). The asterisk indicates BSA in the interaction buffer. (**e**) Zoom-in into interface II of the BiP-GRP94 loading complex. Interface II of the cytosolic loading complex is shown for comparison. (**f**) Ni-NTA pull-down analysis of His_6_-tagged BiP and Strep-tagged GRP94 Δ72 interface II mutant R116G and quantification of the pull-down experiment (n = 3, mean +/- sd). The asterisk indicates BSA in the interaction buffer.

Our model of complex II matches the overall topology of the cytosolic loading complex (**Extended Data Figure 5c**), however shows slightly modified interaction interfaces between the two BiP and GRP94 proteins. At the heart of the complex is the semi-closed conformation of GRP94, supported by the rotation of the GRP94_NTD_s compared to the pre-loading complex (**Extended Data Video 1**). This conformational change increases the distance between the GRP94_MD_ and both the ATP lid and the central GRP94_NTD_ helix, yet puts N-terminal residues K95/K97 in close proximity, a conformation which is supported by the XL-MS data shown in **Figure 3c**. The semi-closed GRP94 conformation is stabilized by the binding of two BiP molecules. Specifically, in both GRP94 protomers the tip of the central GRP94_NTD_ helix (residues 98-121) faces the BiP_NBD_, establishing an interaction with lobe IB (**Figure 5e** and **Extended Data Figure 6b**). Notably, this interaction is dependent on the rotation of the GRP94_NTD_s (**Extended Data Video 1**). The resulting interface II from the GRP94_NTD_-BiP-1 interaction resembles the cytosolic loading complex (**Figure 5e**). Comparison of the paralogs indicates conserved hydrophobic contacts with the C-terminal part of the GRP94_NTD_ helix (residues 117-121) and polar contacts with R116 (**Figure 5e**). Our model indicates variances at interface II’ between GRP94_NTD_ and BiP-2 (**Extended Data Figure 6b**). Rather than packing against the alpha-helical part of lobe IB, the GRP94_NTD_ helix engages adjacent loop regions in lobe IB. This asymmetry in the GRP94-BiP-1/BiP-2 interfaces may be introduced by the more pronounced interaction of BiP-2 with the GRP94 Δ72 purification tag.

To evaluate the importance of the conserved interface II, shown in **Figure 5e**, we analyzed a GRP94 point mutation R116G (R60G in cytosolic Hsp90, R46G in yeast Hsp90), previously shown to decrease the interaction between cytosolic Hsp70/90 proteins and to reduce the fitness of yeast cells^24^. Pull-down experiments showed a ∼30% decrease in the ability of this mutant to bind BiP (**Figure 5f**), corroborating the importance of the conserved interface II for the stability of a BiP-GRP94 Δ72 complex. The comparatively small reduction in the bound GRP94 Δ72 mutant protein can be attributed to the fact that interface I is critical for the pre-loading and loading complex conformation, while the role of interface II is limited to the stabilization of the loading complex. (**Figure 4b**). Overall, our data suggest that through interactions along interface I, BiP establishes the initial contact with GRP94, which is still in an open conformation. Single point mutations at interface I strongly impair the interaction between BiP and GRP94. The binding of a second BiP molecule via interface I and the following formation of interface II stabilize the GRP94 semi-closed state.

## Discussion

GRP94 is a structurally dynamic molecular chaperone critical for the folding, maturation, and degradation of important secretory and transmembrane proteins^19^. In the apo or ADP-bound state, the GRP94 dimer resides in an open conformation^31,32,36^. To tightly interact with its substrate proteins, the dimer closes^32^. Cytosolic Hsp90 proteins rely on co-chaperones for controlled ATP-driven transitioning from the open to the closed state^2,30^. In the secretory pathway, the Hsp70-protein BiP regulates GRP94 function^36,37,42^. Extensive characterization of Hsp90 proteins shows that closure intermediate states play a critical role in the chaperone cycle^24,65–67^ and that BiP is critical for the stabilization of such GRP94 states^36^.

Building on this previous work, our structural and biochemical data provide insight into the BiP-GRP94 *pre-loading* and *loading complex* conformations, two successive states along the GRP94 chaperone cycle (**Figure 6**). These transition states likely became visible by an interaction of BiP with the N-terminal purification tag of GRP94 Δ72 acting as a BiP substrate mimetic and possibly preventing the full closure of the GRP94_NTD_s. In the context of the stabilized complex, we could characterize interactions between the BiP_NBD_ and GRP94. Based on our findings, we propose a model according to which the step-wise binding of BiP controls GRP94 dimer closure. Initially, one BiP molecule binds to the open twisted V conformation of GRP94 by establishing interactions at interface I. The residues critical to this interaction site in the BiP_NBD_ and GRP94_MD_ are highly conserved from bacteria to the different Hsp70/90 chaperone systems in mammalian cells ^24–26,28,37^, indicating that the formation of the pre-loading complex may also be conserved. Following the initial contact established along interface I, additional interactions between BiP and GRP94 are formed along interface II, favoring the loading complex conformation. Importantly, in the loading complex, a second BiP molecule is bound to GRP94 protomer B, increasing the interface area of the BiP-GRP94 complex. We propose that the simultaneous engagement of both GRP94 protomers in interface II promotes GRP94_NTD_ rotation (**Figure 6**). Upon rotation, the GRP94_NTD_s are poised to dimerize and assume the semi-closed conformation. The interactions along interface II thereby define the semi-closed GRP94 state and may underlie the GRP94 closure-accelerating activity of BiP^36,37^.

**Figure 6.**
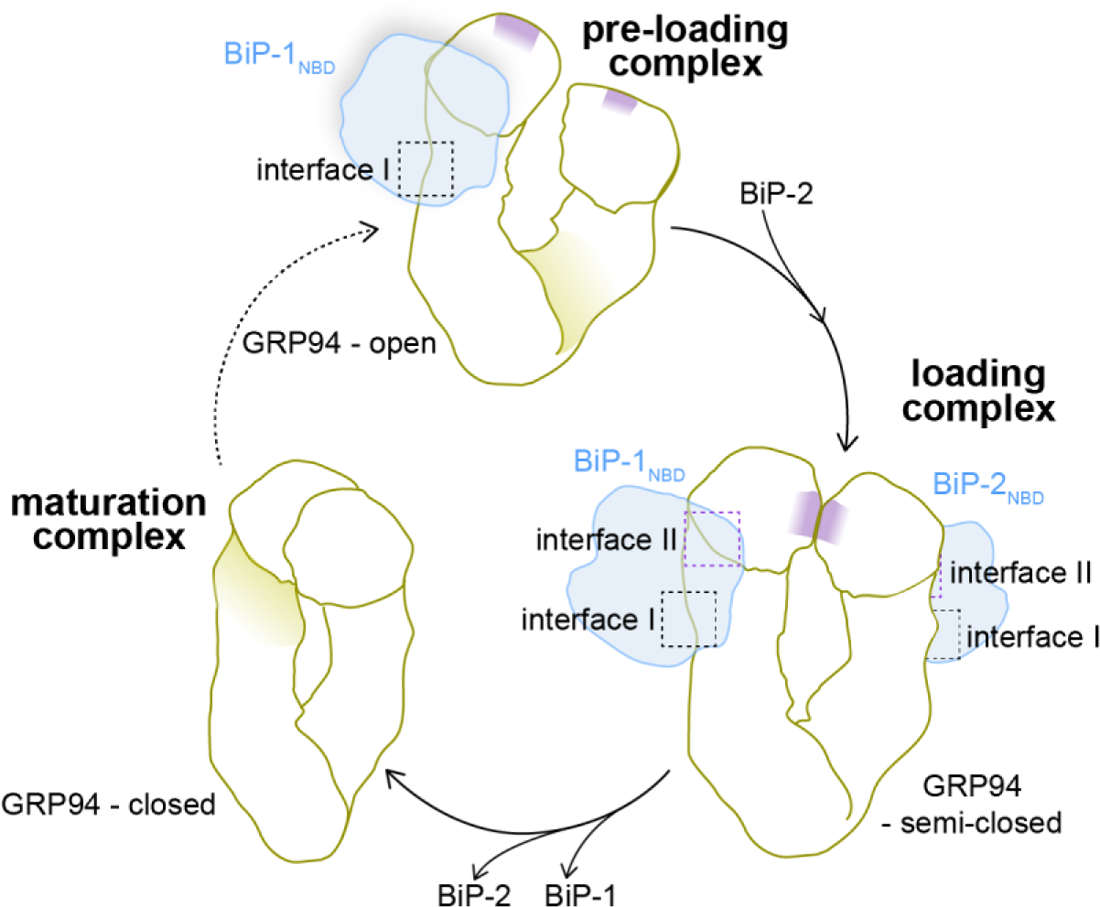
Model of the conformational plasticity of the BiP-GRP94 complex along the GRP94 chaperone cycle. In the pre-loading complex one BiP molecule is bound to an open GRP94 dimer. The binding of a second BiP molecule supports transitioning into the loading complex, where GRP94 is in a semi-closed conformation. Finally, the GRP94 dimer transitions into the fully closed state and BiP concurrently dissociates. The relative position of the GRP94_NTD_ semi-closed dimer interface is indicated in purple.

The cytosolic loading complex is stabilized by the co-chaperone Hop through direct contacts with all complex components – Hsp70, Hsp90, and the substrate protein. Specifically, the semi-closed Hsp90 conformation is defined by direct Hop-Hsp90 interactions and by the binding of two Hsp70 proteins^24^. Even in the absence of a bridging co-chaperone like Hop, two BiP proteins are important for stabilizing the semi-closed GRP94 conformation. The additional role of Hop in substrate binding may not be required for secretory and transmembrane protein folding or may be substituted by still uncharacterized co-chaperones. Given the similarities between the BiP-GRP94 and cytosolic Hsp70-Hsp90 complexes, and the conservation of interface residues, we believe that the stepwise BiP/Hsp70 binding model to drive GRP94/Hsp90 dimer closure (**Figure 6**) may also apply to more sophisticated systems, where co-chaperones have evolved to support and fine-tune this principal mechanism of Hsp70/90 cooperation. Consistent with previous work in the Hsp90 field^24^, our data show that the ATP-bound, fully closed GRP94 does not interact with BiP (**Figure 1d** and **Extended Data Figure 1c**). Closing of the GRP94_NTD_ ATP lid breaks contacts at interface II, subsequent swapping of the GRP94_NTD_s and ATP binding to BiP leaves substrate-bound GRP94 and BiP in the open, domain-docked conformation primed for another round of substrate recruitment (**Figure 6**). Agard and colleagues also proposed a step-wise release of Hsp70-proteins during the transition from the loading complex to the so-called maturation complex, which may equally apply to the simpler BiP-GRP94 system^24,68^ (**Figure 6**).

The coordinated interactions between BiP and GRP94 are aimed at transferring substrates from the BiP_SBD_ to GRP94. We applied the chemical biology tool hydrophobic tagging to translate chaperone-substrate interactions previously identified in cells^46,47^ into an *in vitro* reconstituted system. This shows that hydrophobic tagging provides an efficient approach for *in vitro* reconstitution of chaperone-substrate complexes as demonstrated by our usage for two different types of chaperones, BiP and GRP94 (**Figure 2**). Inducing HT2 misfolding in the presence of chaperones may provide an advantage in establishing chaperone-substrate complexes over an experimental setup where protein misfolding and chaperone-binding reactions are separated. BiP and GRP94 individually interacted with HT2^misfolded^ but could also form a multi-chaperone-substrate complex. While we could not visualize this complex, we expect, by extrapolating from the cytosolic loading complex, a state where the substrate protein is engaged by BiP and GRP94 simultaneously. We presume that interactions between substrate protein and GRP94 drive the formation of the loading complex, in addition to interactions between the GRP94_NTD_s and BiP_NBD_s, further facilitating substrate transfer. The orientation of the BiP_SBD_ with respect to the BiP_NBD_ captured in our sample underlines the flexibility of the BiP_SBD_ in the domain-undocked state. In-solution studies have previously highlighted that the crystal structures of the domain-docked and domain-undocked BiP conformations (**Figure 1a**) represent endpoint states^39,40^. This flexibility of the BiP_SBD_ will facilitate the delivery of a highly diverse range of substrate proteins to GRP94.

Our negative-stain EM and XL-MS data highlight striking similarities between the comparatively simple BiP-GRP94 chaperone system and their intricate Hsp70/90 paralogs and point to a highly conserved mode of substrate transfer among Hsp70/90 systems. Thereby Hsp70/BiP proteins play a conserved role, not only in the delivery of substrate proteins but also in the activation of Hsp90/GRP94 proteins. On the basis of the identified BiP-GRP94 pre-loading and loading complexes, it will be interesting to define ER-specific regulatory mechanisms. Post-translational modifications may influence transitioning between the different states of the GRP94 chaperone cycle. Regulatory modifications may affect the stability and lifetime of the different BiP-GRP94 complex states and could also change the role of the chaperones from collaborating in protein folding to supporting protein degradation.

## Supporting information

Extended Data Figures (combined)

## Acknowledgements

We would like to thank Marie-Anne Derichs, Daniel Agranovski, and Leonhard Sewald for supporting biochemical experiments. This work was funded by the Sofja Kovaleveskaja Award by the Alexander von Humboldt Foundation endowed by the Federal Ministry of Education and Research. D.H., F.K., M.K., S.P. and E.S.-G. were supported by the Deutsche Forschungsgemeinschaft (DFG, German Research Foundation) – SFB1430 – Project-ID 424228829. E.S.-G. further acknowledges instrumentation funding under the Großgerätinitiative - Project number: 436586093 and the support of Germany’s Excellence Strategy - EXC-2033 - Project number: 390677874. Screening and data acquisition of EM data were performed at the StruBiTEM Facility of the University of Cologne, with support by Monika Gunkel and Elmar Behrmann.

## Author contributions

J.C.B. conducted all biochemical experiments and prepared MS samples. F.K. and M.K. processed MS samples and analyzed the data together with J.C.B. J.C.B and L.Z. prepared samples for EM. L.Z. and S.P. performed EM data collection, processing, and analysis. Y.A.H. carried out the structural modeling. Y.A.H. and E.S.G. designed, analyzed, and wrote the results and discussion of the computational modeling. D.H. outlined the project and prepared the manuscript with input from all co-authors.

## Methods

### Reagents and antibodies

HyT36 and HyT36(-Cl) were a kind gift from Craig Crews and George Burslem ^50,51^. 6-Chlorohexanol (Sigma-Aldrich, C45008), Trypsin (Sigma-Aldrich, T8003), DSBU (ThermoFisher, A35459), Glutaraldehyde (ThermoFisher), ATP (Sigma-Aldrich, A6419), AMP-PNP (Sigma-Aldrich, A2647), Anti-His5 (Santa Cruz Biotechnologies, SC-8036), Anti-Strep (Qiagen, 34850), Goat anti-mouse HRP-coupled secondary antibody (Invitrogen, 31444) were from commercial suppliers.

### Plasmids and molecular cloning

The expression plasmids His_6_-linker-TEVsite-GRP94 (73-802) in pET151 and His_6_-BiP (27-655) in pNIC28-Bsa4 were a kind gift from Timothy Street^37^. GRP94 was re-cloned into pET21d to yield an expression plasmid encoding Strep-linker-TEVsite-GRP94 (73-802). See **Extended Data Figure 6a** for details on the N-terminal purification tag. Strep-GRP94 FL was cloned from C2C12 cDNA into pET21d by Gibson assembly (NEB Hifi Builder). His_6_-BiP-NBD (27-411) was cloned from His_6_-BiP in pNIC28-Bsa4 via Gibson assembly into the same vector. pcDNA5/FRT/TO-EGFP-HT2^46^ was used as a template to clone HT2-His_6_ in pET21a. Spot-HT2 was cloned from His_6_-HT2 into pET22b-SUMO. All point mutations were introduced by site-directed mutagenesis. All primers are listed in the table below.

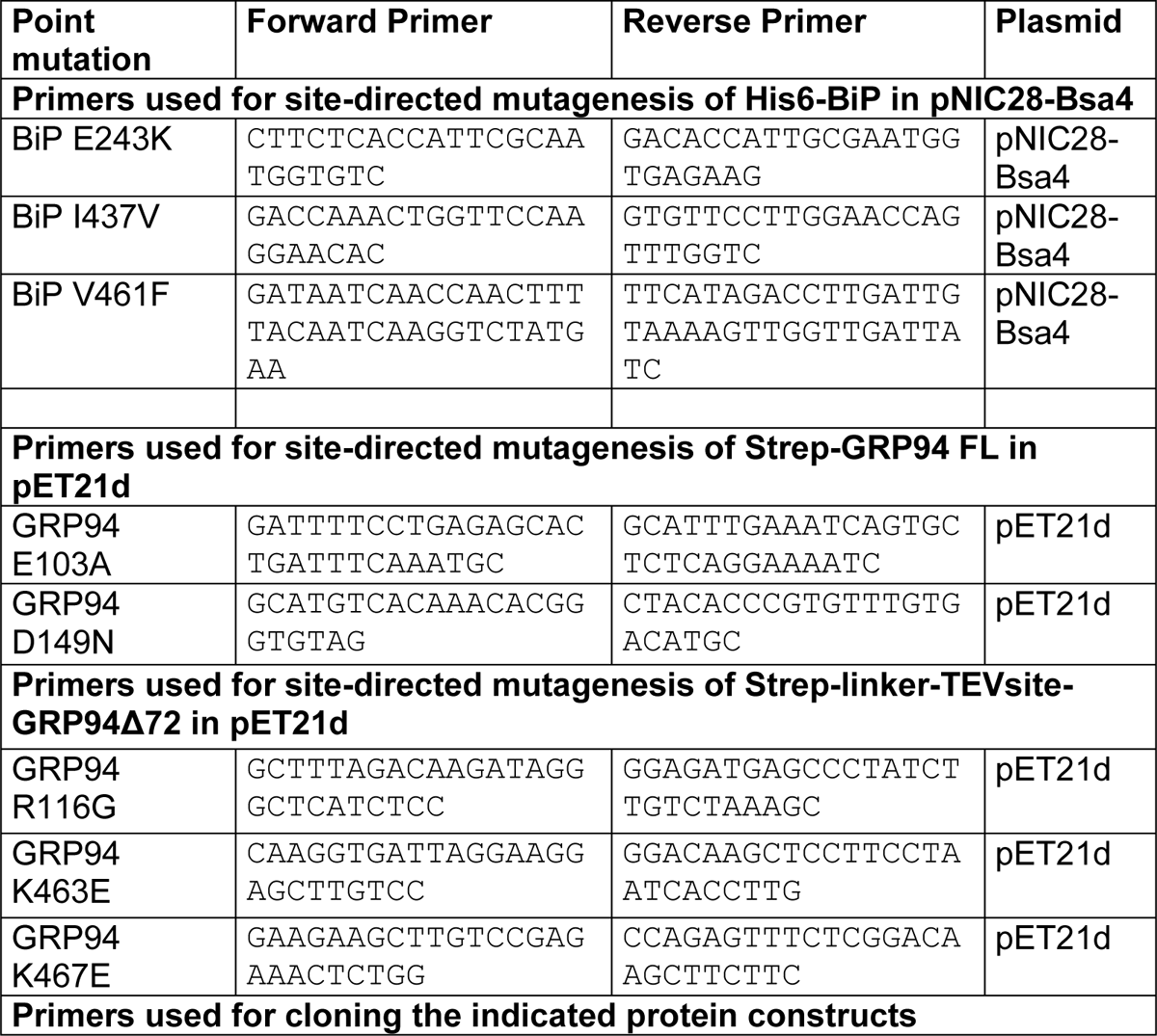
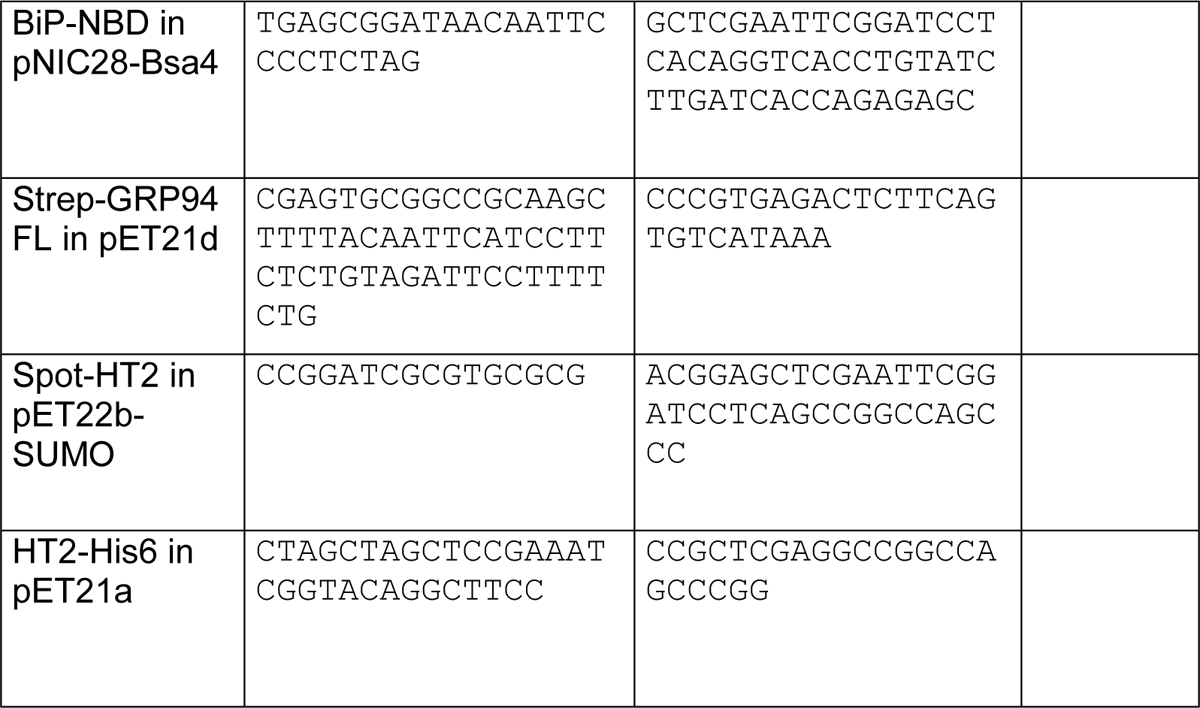

### Protein expression and purification

All proteins were expressed in *Escherichia coli* BL21(DE3) cells. Protein expression was induced with 250 μM isopropyl β-D-1-thiogalactopyranoside (IPTG) for 16 h at 18 °C for GRP94 and BiP proteins and for 5 h at 25 °C for His_6_-SUMO-HT2 and HT2-His_6_. Cells were harvested by centrifugation, resuspended in 50 mM Na_2_HPO_4_ pH 8.0, 300 mM NaCl for His_6_-SUMO-HT2 and HT2-His_6_, 100 mM HEPES pH 8.0, 400 mM NaCl, 10 mM imidazole, 0.5 mM tris(carboxyethyl)phosphine (TCEP) for His_6_-BiP proteins and 100 mM HEPES pH 8.0, 400 mM NaCl, 0.5 mM TCEP for Strep-tagged GRP94 proteins and lysed by sonication.

For His_6_-tagged proteins, cleared cell lysate was applied to a HisTrap HP column (Cytiva). The column was washed by applying a stepwise imidazole gradient and the His_6_-tagged protein was eluted in 150 mM imidazole in lysis buffer. His_6_-SUMO-Spot-HT2 was digested with His_6_-Ulp1 and subjected to another round of HisTrap affinity chromatography to isolate Spot-HT2. For Strep-tagged proteins, cleared cell lysate was applied to a StrepTrap HP column (Cytiva) and eluted with 2.5 mM D-desthiobiotin in lysis buffer. As a final step all proteins were subjected to size-exclusion chromatography using a Superdex 200 16/600 column equilibrated with 50 mM HEPES pH 7.5, 150 mM NaCl for the HT2 proteins and with additional 0.5 mM TCEP for BiP and GRP94 proteins. Fractions containing the protein of interest were concentrated, flash frozen in liquid N_2_ and stored at −70 °C.

### Hydrophobic tagging

Spot-HT2 or HT2-His_6_ was incubated with HyT36, HyT36(-Cl), or dimethyl sulfoxide (DMSO) for 10 min at RT. To induce misfolding samples were incubated at 37 °C.

### Limited proteolysis

10 µM His_6_-HT2 reacted with 12 µM HyT36 or in presence of 12 µM HyT36(-Cl) or DMSO was incubated with 0.1 µM trypsin (Sigma-Aldrich) for 15, 30, 60, and 90 min at 37 °C. Reactions were stopped by addition of SDS-PAGE sample buffer and resolved on SDS-PAGE gels.

### CD spectroscopy

CD spectra of 10 µM Spot-HT2 reacted with 10 µM HyT36, 10 µM 6-chlorohexanol, or in presence of DMSO were recorded in 20 mM Tris pH 7.5, 100 mM NaCl at 25 °C and 37 °C at a Jasco Spectropolarimeter J-710 connected to the temperature control unit Jasco PFD-350s with a Julabo F-250 compact recirculation cooler at 0.2 nm steps with 10 scans. Data were analyzed in GraphPad Prism. Given the presence of DMSO in the samples, spectra were analyzed from 218 to 260 nm.

### Sample preparation for electron microscopy (EM)

To prepare protein complexes for negative-stain EM, 16 µM BiP and 32 µM HT2 reacted with 32 µM HyT36 were incubated for 30 min at 37 °C in 50 mM HEPES pH 7.5, 50 mM NaCl, 2 mM MgCl_2_, 50 µM ATP. Then 16 µM GRP94 were added to the mix and the sample was incubated for another 30 min at 37 °C. Gradients were generated by using a range from 5% glycerol in 50 mM HEPES, 50 mM NaCl, 1 mM CaCl_2_ to 45% glycerol in 50 mM HEPES, 50 mM NaCl, 1 mM CaCl_2_, 0.5% glutaraldehyde using the Master Gradient 108 (BioComp Instruments, New Brunswick, Canada, program 5-45% Sucrose, 45 s). The generated gradient was incubated for 60 min at 4 °C. The protein samples were carefully applied to the gradient and centrifuged for 16 h at 130,000 g at 4 °C (Optima XPN-80 Ultracentrifugation from Beckman Coulter). 200 µl fractions were taken from the top of the gradient and the glutaraldehyde quenched with 20 mM Tris pH 7.5. Gradient fractions were analyzed by SDS-PAGE.

To generate negative-stain grids 4 µl of the GraFix fraction was applied to a Copper 400 Mesh grids with continuous carbon (Electron Microscopy Sciences) after glow discharge, incubated for 30 s, immediately blotted with filter paper and stained via five successive short incubations of 2% (w/v) uranyl formate (Science Services). The excess stain was removed with filter paper and the grids dried before imaging.

### Collection and processing of negative-stain EM data

Grids were imaged using a Talos L120C microscope (Thermo Fisher) operated at 120 keV with a CMOS camera (Ceta 16M). Micrographs were acquired at a nominal pixel size of 1.89 Å and 25 e^-^/Å^2^ total dose. Data was processed in CryoSPARC^69^, using CTFFIND4 for CTF estimation^70^, Blob picker for particle picking and iterative rounds of reference-free 2D classification. Classes representing damaged protein or artefacts were removed in several rounds of 2D classification. Approximately ∼434,000 particles corresponding to representative 2D class averages were used for *ab initio* reconstructions with several classes and heterogeneous refinements (see **Extended Data Figure 3b** and 4).

### Molecular modeling

The pre-loading state of the GRP94-BiP complex was modeled with the structures of the “open” ADP-bound GRP94 (PDB ID: 2O1V^31^) and the structure of the nucleotide-binding domain (NBD) of BiP (PDB ID: 6DWS) as inputs. The region comprising residues 393-408 of GRP94 was modelled using Modeller^71,72^. For the loading complex, a homology model of GRP94 was obtained with the SWISS-MODEL web server^73^ using the “semi-closed” structure of HSP90 (PDB ID: 7KW7^24^) as template. This complex encompassed one BiP-NBD chain as in the pre-loading complex and a model of the extended conformation of BiP which also included the substrate-binding domain (SBD, residues 30-635). This structure was built using SWISS-MODEL with the *E. coli* HSP70 homolog DnaK (PDB ID: 2KHO^64^) as template. For this model, the region of the residues 285-330 was truncated similar to the open structure.

Before fitting into the EM density, the α-helical C-terminal region of the model of the extended structure of BiP was truncated (residues 530-635 were removed). For the modeling based on the obtained negative-stain EM maps, a first rigid body fitting was performed with UCSF Chimera^74^ on the densities obtained for the pre-loading and loading complexes. Subsequently, a refinement of the fitting was done with Rosetta’s *relax* application ^62^ with the specific flags for cryo-EM refinement^63^. We used the cross-linking information corresponding to NBD-NBD, SBD-SBD, and NBD-SBD pairs as constraints for the fitting. For all the fittings the input resolution of the density maps was 20 Å. For both pre-loading and loading complexes, a cartesian fitting was employed and 5000 decoys were generated. For the evaluation, the Rosetta’s *ref2015*^75,76^ and *elec_dens_fast* score functions were employed, the latter with a weight = 10. Additionally, the real-space correlation coefficient (RSCC) was calculated with Rosetta’s *density_tools* application. The interface area was measured with UCSF ChimeraX^77^. UCSF Chimera^77^ and Pymol (Molecular Graphics System, Version 2.0 Schrödinger, LLC) were employed for the analysis and visualization of the structures.

### Sample preparation for XL-MS

All reactions were carried out in 50 mM HEPES pH 7.5, 50 mM NaCl, 2 mM MgCl_2_. For generating the BiP-GRP94 complex 4 µM BiP and 4 µM GRP94 Δ72 were incubated for 60 min at 37 °C. The trimeric complex of BiP, GRP94 Δ72 and HT2 was generated via reaction of 8 µM HT2 with 12 µM HyT36 for 10 min at RT. 4 µM BiP was added and the sample incubated for 30 min at 37 °C. 4 µM GRP94 Δ72 was then added and incubated for an additional 30 min. Samples were brought to RT (10 min) and disuccinimidyl dibutyricurea (DSBU, ThermoFisher, A35459) was added to a final concentration of 0.5 mM. Crosslinking reactions were incubated for 20 min at RT, quenched with 20 mM Tris pH 7.5, and the protein complexes isolated by SDS-PAGE.

### In-gel digestion (IGD) of DSBU-crosslinked proteins

Gel regions containing crosslinked proteins were excised and washed 2× with water and 2× with 100 mM ammonium bicarbonate solution. The proteins were reduced with 10 mM TCEP (30 min, 62 °C) while gently shaking and subsequently alkylated by adding 55 mM iodoacetamide (IAA) (30 min, room temperature, in the dark). The gel slabs were subsequently washed 3× with acetonitrile (3× 50 µl). After the last wash, the gel slabs were completely dried by using a vacuum concentrator (Eppendorf) for 5 min. The dried gel pieces were next incubated with 200 µl 10 ng µl^-1^ trypsin (37 °C, 16 h, vigorous shaking). The digestion was stopped by addition of formic acid to a final concentration of 5% (v/v) formic acid. The digestion supernatant was transferred to a fresh Eppendorf tube. The remaining gel pieces were subsequently washed 3× with acetonitrile (3× 50 µl). The supernatants of these washes were combined with the recovered digestion mix and dried in a vacuum concentrator (Eppendorf).

The cleared tryptic digests were then desalted on home-made C18 StageTips as described^78^. Briefly, peptides were immobilized and washed on a 2 disc C18 StageTip. After elution from the StageTips, samples were dried using a vacuum concentrator (Eppendorf) and the peptides were taken up in 0.1% formic acid solution (10 μl) and directly used for LC-MS/MS experiments (see below for details).

### LC-MS/MS settings

LC-MS/MS settings. MS Experiments were performed on an Orbitrap LUMOS instrument (Thermo) coupled to an EASY-nLC 1200 ultra-performance liquid chromatography (UPLC) system (Thermo). The UPLC was operated in the one-column mode. The analytical column was a fused silica capillary (75 µm × 28 cm) with an integrated fritted emitter (CoAnn Technologies) packed in-house with Kinetex 1.7 µm core shell beads (Phenomenex). The analytical column was encased by a column oven (Sonation PRSO-V2) and attached to a nanospray flex ion source (Thermo). The column oven temperature was set to 50 °C during sample loading and data acquisition. The LC was equipped with two mobile phases: solvent A (0.2% FA, 2% Acetonitrile, ACN, 97.8% H_2_O) and solvent B (0.2% FA, 80% ACN, 19.8% H_2_O).

All solvents were of UPLC grade (Honeywell). Peptides were directly loaded onto the analytical column with a maximum flow rate that would not exceed the set pressure limit of 980 bar (usually around 0.4 – 0.6 µl min^-1^). Peptides were subsequently separated on the analytical column by running a 105 min gradient of solvent A and solvent B (start with 2% B; gradient 2% to 8% B for 7:00 min; gradient 8% to 22% B for 60:00 min; gradient 22% to 40% B for 25:00 min; 40% to 98% B for 3:00 min; 98% for 10 min) at a flow rate of 350 nl min^-1^. The mass spectrometer was controlled by the Orbitrap Fusion Lumos Tune Application (version 3.3.2782.28) and operated using the Xcalibur software (version 4.3.73.11). The mass spectrometer was set in the positive ion mode. The ionization potential (spray voltage) was set to 2.1 kV. Source fragmentation was turned off. Precursor ion scanning was performed in the Orbitrap analyzer (FT; fourier transform mass spectrometer) in the scan range of *m/z* 370-1600 and at a resolution of 120000 with the internal lock mass option turned on (lock mass was 445.120025 *m/z*, polysiloxane) ^79^. AGC (automatic gain control) was set to “standard” and acquisition time to “auto”. Product ion spectra were recorded in a data dependent fashion in the Orbitrap at a variable scan range (“auto”) and at a resolution of 15000. Peptides were analyzed using a “top speed” regime (repeating cycle of full precursor ion scan (AGC target “standard” acquisition time “200 ms”) followed by dependent MS2 scans for 3 seconds (minimum intensity threshold 2×104)). The MS2 precursor ions were isolated using the quadrupole (isolation window 2 m/z) and fragmentation was achieved by Higher-energy C-trap dissociation (HCD) (normalized collision mode set to “stepped” and normalized collision energy set to “27, 30, 33%”). During MS2 data acquisition dynamic ion exclusion was set to 60 seconds. Only charge states between 3-7 were considered for fragmentation.

### XL-MS data analysis

Thermo RAW files were converted to the mzML or mfg file format using ProteoWizard (version: 3.0.23018-60066e9)^80^ with default settings and submitted to Xisearch (version: 1.7.6.7) for further analysis^81^. For each search a dedicated database containing only the sequences of the investigated proteins was used. The following settings were applied: MS1 error tolerances of 6 ppm; MS2 error tolerances of 20 ppm; trypsin digestion with 2 missed cleavages allowed.

Fixed modifications: carbamidomethylation on cysteine. Variable modifications: oxidation on methionine. Crosslinker: DSBU, linked amino acids are lysine, serine, threonine, tyrosine, linear modifications NH_2_, OH, Bu, BuUr; max modification per peptide: 3. Ions: fragment B-Ion and Y-Ion; The resulting candidates from Xisearch were filtered to 5% false discovery rate (FDR) on residue pair level using the software xiFDR (version 2.1.5.5)^82^. The mgf-files, xiFDR results and FASTA sequences were then imported and visualized by XiView to generate Figures 3a and Extended Data Figure 3a^83^.

### Pulldowns and sequential pull-down

All reactions were carried out in 50 mM HEPES pH 7.5, 50 mM NaCl, 2 mM MgCl_2_, 50 µM ATP, 3 µM BSA. 8 µM HT2 reacted with 12 µM HyT36 was incubated with 4 µM His_6_-tagged BiP protein, 4 µM Strep-tagged GRP94 protein or both (as indicated in the individual figures) for 60 min at 37 °C. For pull-downs involving untagged GRP94 Δ72, the purification tag was removed by incubation with His-GST-TEV protease for 2h at RT in 50 mM HEPES pH 7.5, 50 mM NaCl, 1 mM CaCl_2_. Upon protease cleavage, untagged GRP94 Δ72 protein was isolated from the flow-through of a Streptactin and following Ni-NTA purification. 4 µM His_6_-tagged BiP protein and 4 µM Strep-tagged or untagged GRP94 protein were incubated for 60 min at 37 °C.

For His_6_-tagged BiP protein pull-down, samples were incubated with Ni-NTA beads (Qiagen) for 60 min at 37 °C, washed with 3x with 50 mM HEPES pH 7.5, 50 mM NaCl, 10 mM imidazole, 1 mM CaCl_2_ and finally eluted with 500 mM imidazole in wash buffer. For Strep-tagged GRP94 protein pull-down, samples were applied to StrepTactin 4 flowXT gravity flow columns (IBA), washed 3x with 50 mM HEPES pH 7.5, 50 mM NaCl, 1 mM CaCl_2_ and finally eluted with 50 mM biotin in wash buffer. For the sequential pull-down, the elution of a Strep-pull-down (as described above) was used as the input for a Ni-NTA pull-down (as described above). All input and elution fractions were applied to SDS-PAGE. Quantifications were performed in ImageLab (BioRad) and data were plotted in Prism (Graphpad).

### Analytical SEC

BiP and GRP94 proteins were incubated at a final concentration of 8 μM (if not otherwise indicated) for 60 min at 37 °C and applied to a Superose 6 PC 3.2/300 column in 50 mM HEPES pH 7.5, 50 mM NaCl, 1 mM CaCl_2_ on an Ettan LC system. Indicated fractions were analyzed by SDS-PAGE.

### Glutaraldehyde crosslinking

For glutaraldehyde crosslinking experiments, 8 µM GRP94 proteins and 8 µM BiP_NBD_ were incubated separately or together in 50 mM HEPES pH 7.5, 50 mM NaCl, 2 mM MgCl_2_ in presence or absence of 1 mM AMP-PNP for 60 min at 37 °C. Samples were brought to RT (10 min) and 0.006% glutaraldehyde was added. After 5 min at RT, reactions were quenched with 20 mM Tris pH 7.5 and the samples were analyzed by SDS-PAGE and western blot using anti-His_5_ (Santa Cruz Biotechnologies, SC-8036, 1:1000 dilution) and anti-Strep (Qiagen, 34850, 1:1000 dilution) antibodies. Bands were detected by chemiluminescence on a BioRad ChemiDoc imager.

