## Extended Data Figures (combined) for "Conformational plasticity of a BiP-GRP94 chaperone complex"

**CONTENT**

**Extended Data Figures 1-6, Extended Data Table 1**

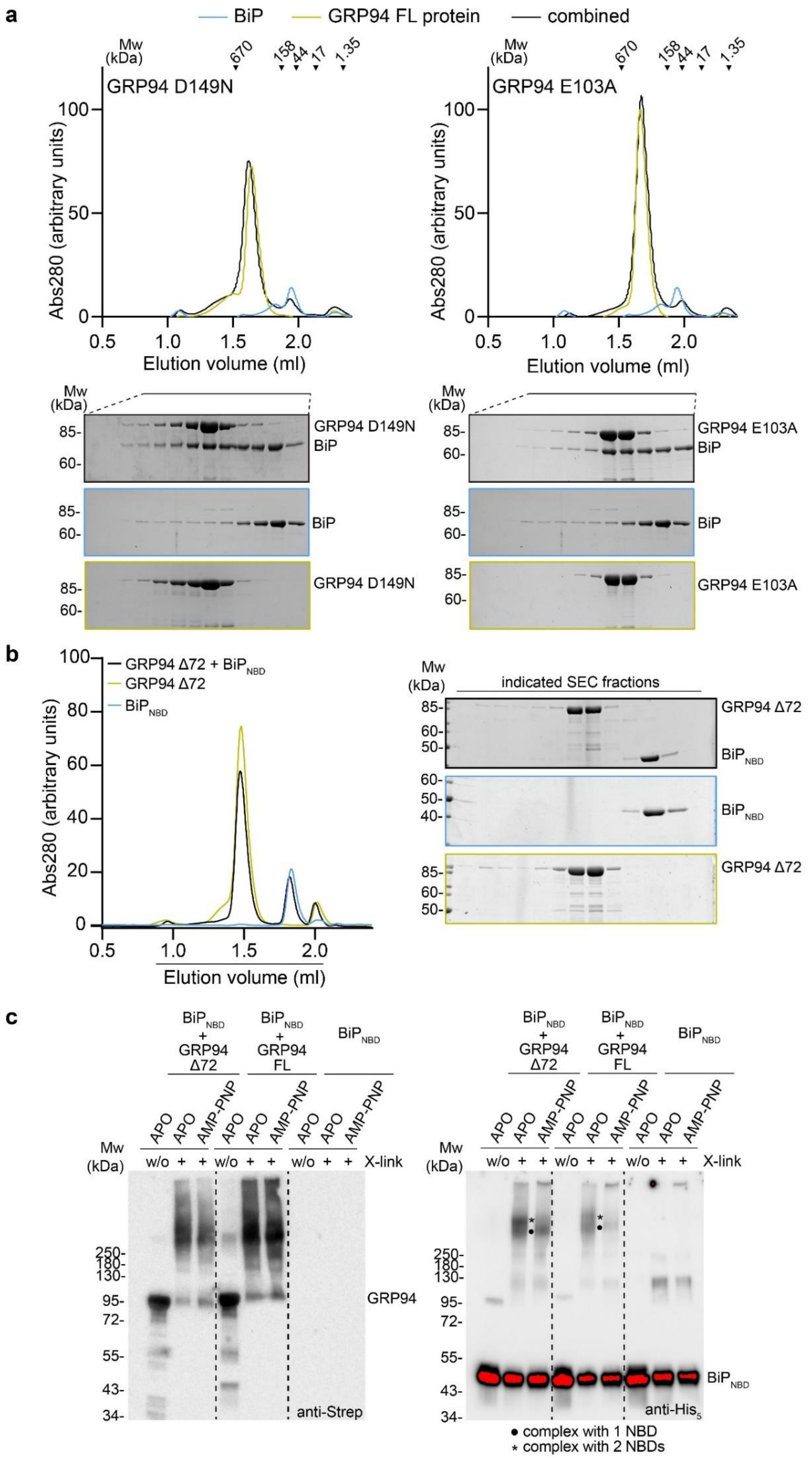
a
BiP
GRP94 FL protein
combined
Mw (kDa)
670
158
44
17
1.35
GRP94 D149N
GRP94 E103A
Abs280 (arbitrary units)
Elution volume (ml)
GRP94 D149N
BiP
GRP94 E103A
b
GRP94 Δ72 + BiP$_{NBD}$
GRP94 Δ72
BiP$_{NBD}$
indicated SEC fractions
c
BiP$_{NBD}$ + GRP94 Δ72
BiP$_{NBD}$ + GRP94 FL
BiP$_{NBD}$
APO
AMP-PNP
w/o
X-link
GRP94
anti-Strep
anti-His$_5$
● complex with 1 NBD
* complex with 2 NBDs

**Extended Data Figure 1. Additional biochemical data on the interaction of BiP and GRP94.** (**a**) Analytical SEC and SDS-PAGE analysis of complex formation between GRP94 full-length (FL) ATP-binding mutant D149N, ATP hydrolysis mutant E103A and BiP. (**b**) Analytical SEC and SDS-PAGE analysis of $BiP_{NBD}$-GRP94 Δ72 complex formation. (**c**) Western blot analysis of glutaraldehyde crosslinking reactions to detect Strep-tagged GRP94 Δ72 or -FL protein and $His_6$-tagged $BiP_{NBD}$ (related to Figure 1d).

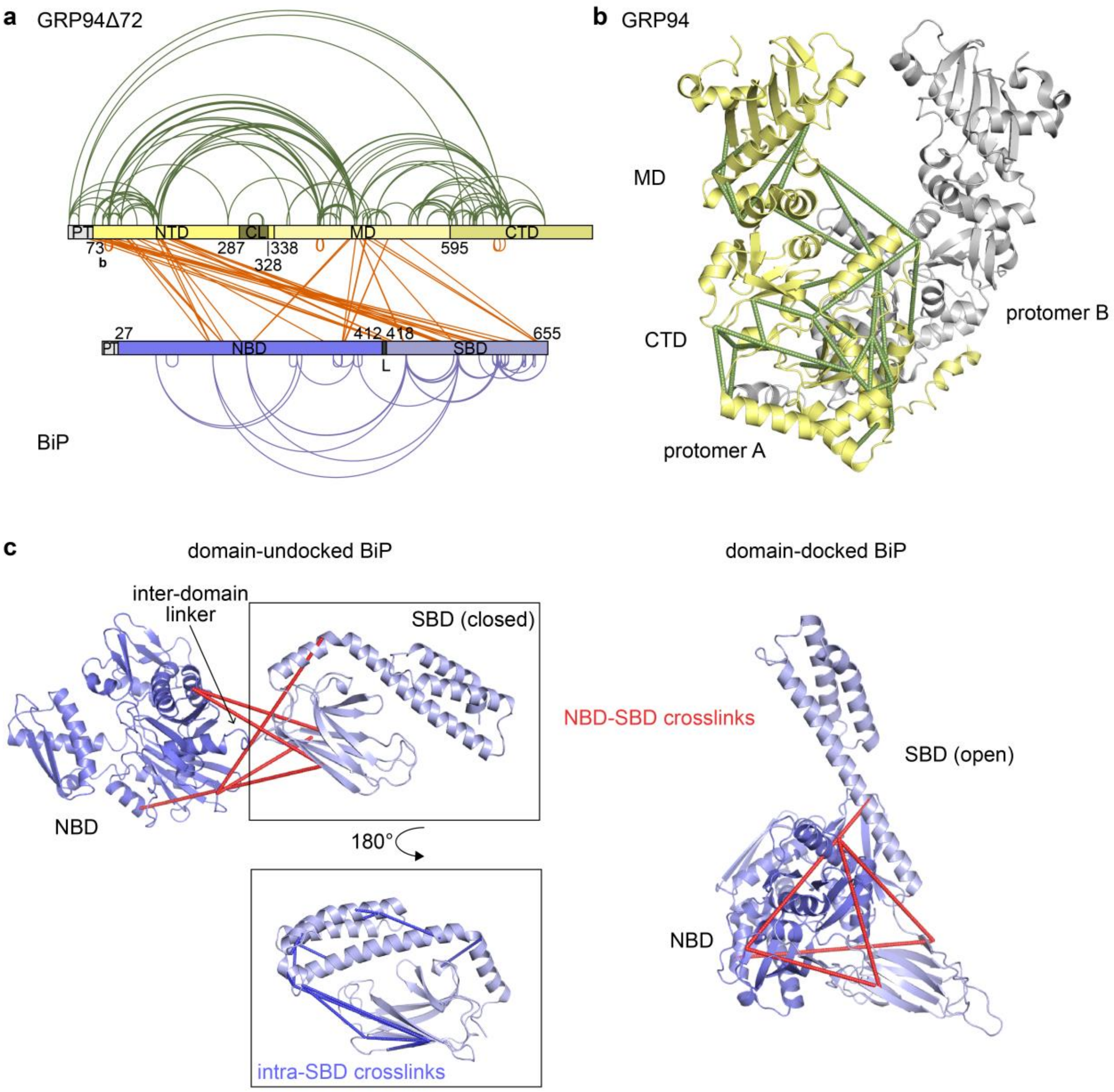


**Extended Data Figure 2. Analysis of XL-MS data.** (**a**) Map of crosslinks identified in the BiP-GRP94 Δ72 sample. (**b**) Intra-GRP94 crosslinks mapped onto a GRP94$_{MD-CTD}$ crystal structure (PDB ID: 2O1T). (**c**) Intra-BiP crosslinks from the BiP$_{NBD}$ to the BiP$_{SBD}$ mapped onto the domain-undocked BiP structure (homology model generated with SWISS-MODEL based on PDB ID: 2KHO) and onto the domain-docked conformation (PDB ID: 5E84). All crosslinks shown in red are > 30 Å. Intra-BiP$_{SBD}$ crosslinks (blue) were mapped onto a closed SBD structure (PDB ID: 5E85) and demonstrate the compatibility with the closed rather than the open SBD conformation.

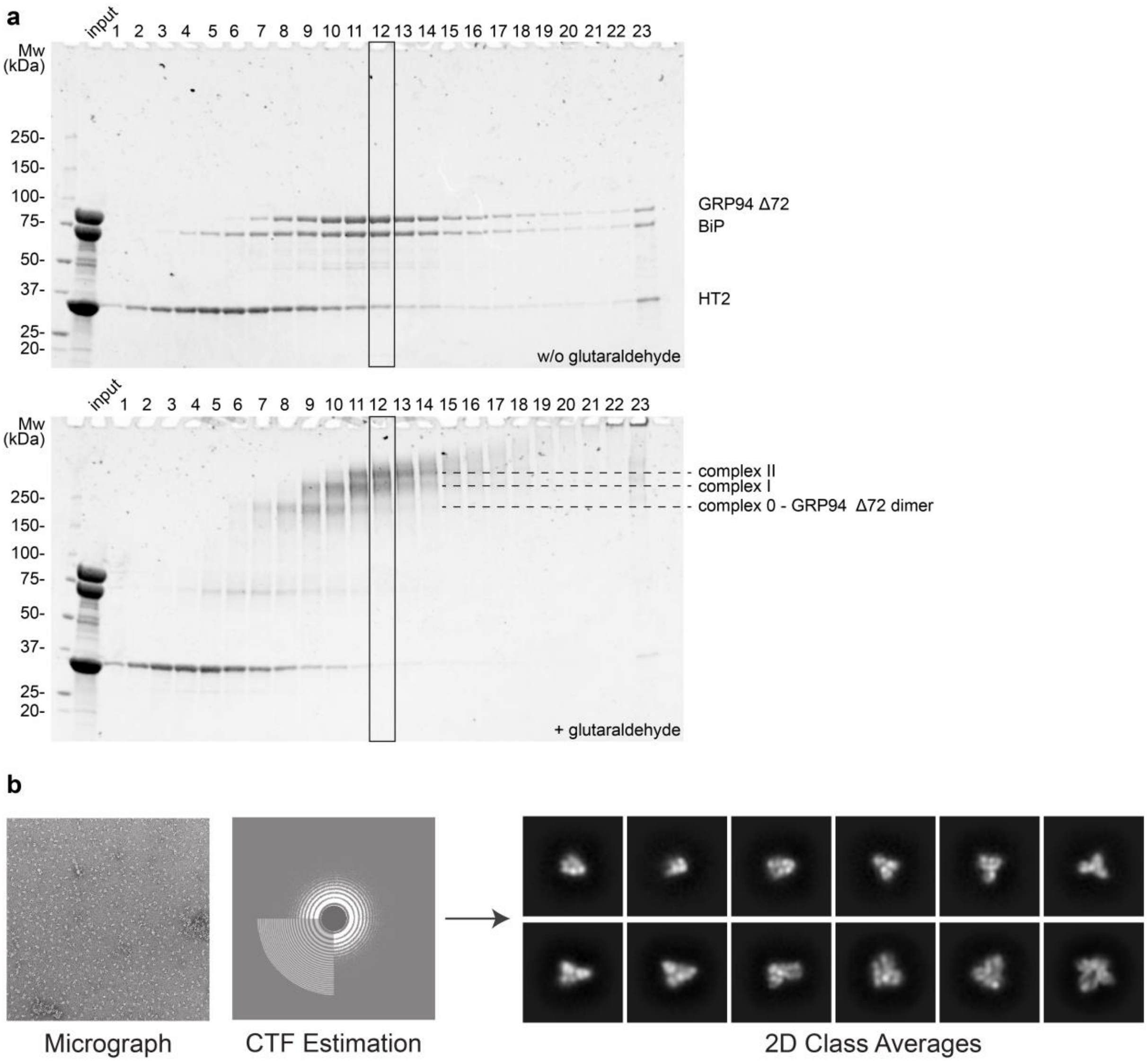


**Extended Data Figure 3. Characterization of negative-stain EM sample.** (**a**) SDS-PAGE gel analysis of GraFix fractions. Gradient density increases with fraction number. Fraction 12 was analyzed by negative-stain EM. (**b**) Representative negative-stain EM micrograph and 2D class averages obtained from fraction 12.

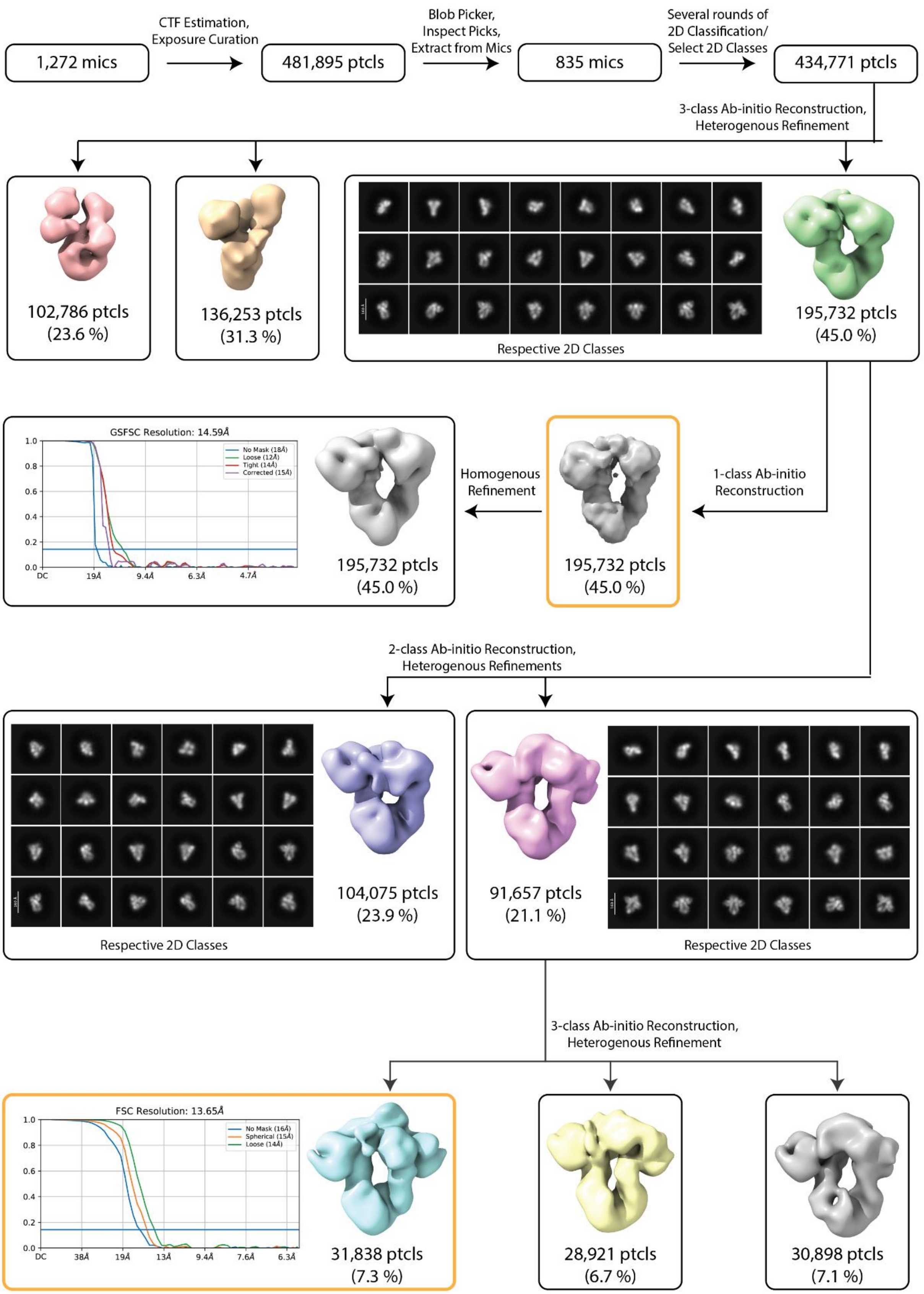


**Extended Data Figure 4. Single particle negative-stain EM image processing pipeline.** Flow chart of classifications and refinements, resulting in reconstructions of the pre-loading and loading complex conformations. Yellow boxes indicate 3D reconstructions that were further used for molecular modeling. The number of particles that went into each reconstruction is indicated, together with their respective percentage share of the initial 434,771 particles. 2D classifications of those particle stacks resulted in the shown respective 2D class averages (scale bar: 160 Å).

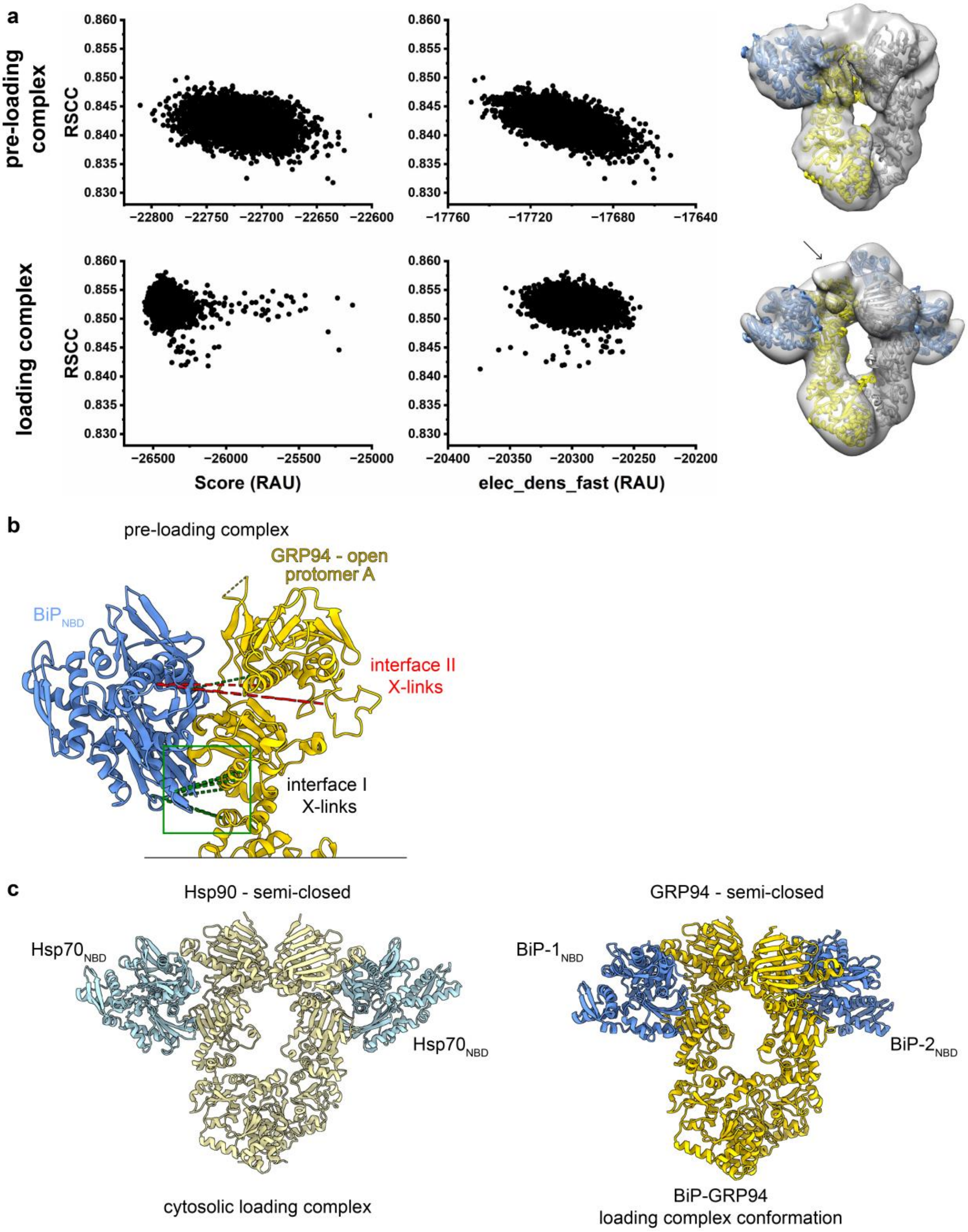


**Extended Data Figure 5. Evaluation of pre-loading and loading complex models.** (**a**) Quality metrics of molecular modeling of the pre-loading and loading complexes. RSCC: Real-space correlation coefficient, RAU: Rosetta arbitrary units. The arrow indicates additional density not filled by the loading complex models. (**b**) Crosslinks identified between BiP$_{NBD}$ and GRP94 Δ72 mapped onto the pre-loading complex model. GRP94 protomer B is omitted for clarity. Red lines indicate crosslinks > 30 Å, green lines < 30 Å. (**c**) Side-by-side comparison of the cytosolic (left) and BiP-GRP94 (right) loading complex conformation. For clarity, exclusively the NBDs of Hsp70/BiP and the Hsp90/GRP94 dimer structures are shown.

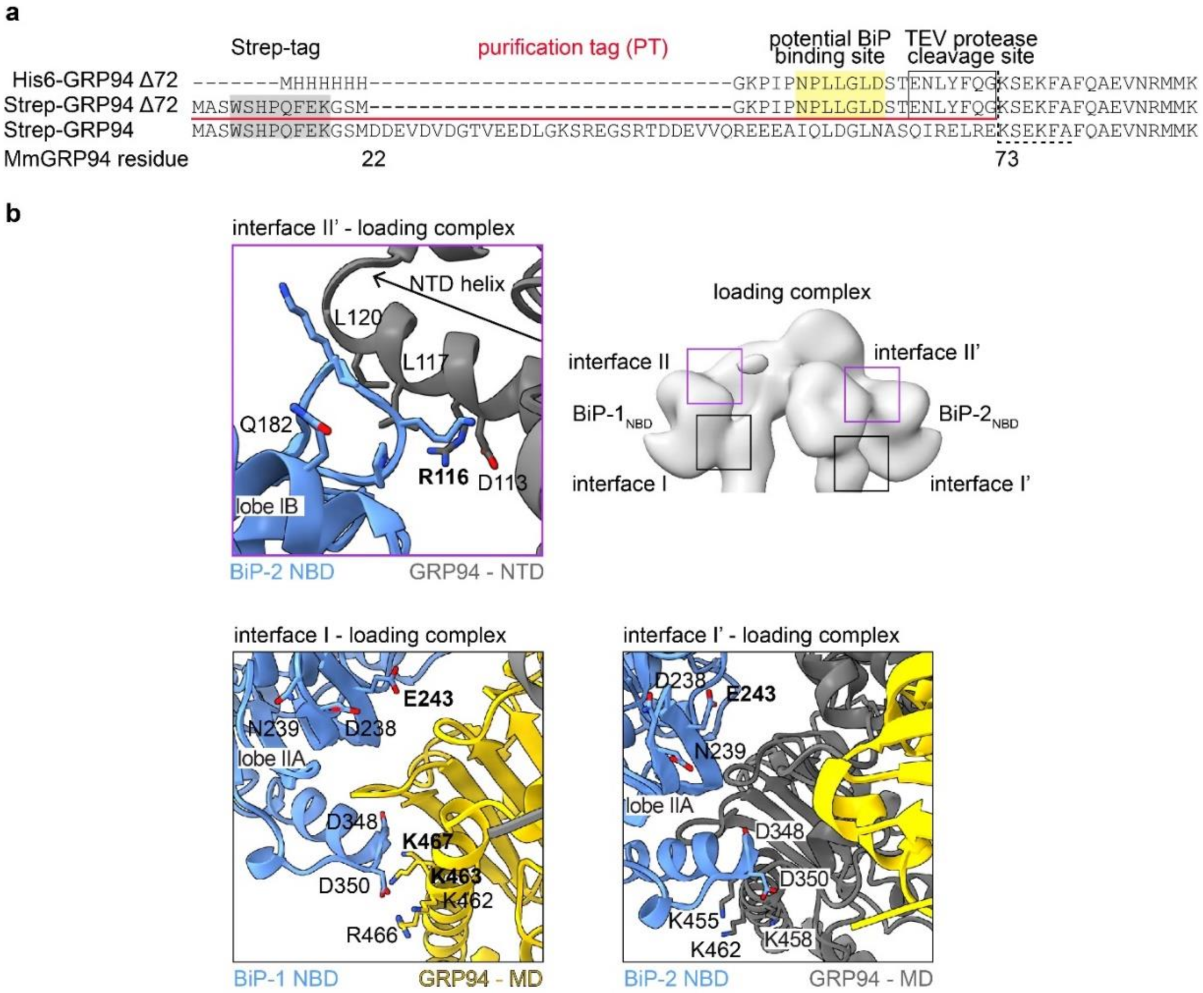


**Extended Data Figure 6. BiP-GRP94 interaction interfaces in the loading complex conformation.** (**a**) Sequence alignment of the indicated expression constructs. A sequence stretch that is part of the $His_6$- and Strep-tagged GRP94 Δ72 constructs highlighted in yellow may serve as a BiP substrate mimetic. (**b**) Zoom-in into interfaces I, I' and II' of the BiP-GRP94 loading complex. Compare to Figure 5b and 5d.

**Extended Data Table 1.** BiP-GRP94 crosslinks identified in the respective samples by XL-MS. Data were analyzed in Xisearch and a score cut-off of 15 was applied. Residues in pink are part of the $BiP_{SBD}$ or $GRP94_{CTD}$, residues in green are part of the $GRP94_{MD}$, residues in blue are part of the $BiP_{NBD}$ or $GRP94_{NTD}$.

**Inter-protein crosslinks of the BiP-GRP94Δ72-HT2 sample**

| BiP residue Uniprot ID P20029 | GRP94 residue Uniprot ID P08113 | Score | Crosslink outlines the following interaction |
|---|---|---|---|
| 164 | purification tag pos. 2 | 18 | purification tag to GRP94 NTD |
| 464 | purification tag pos. 2 | 19 | purification tag to BiP substrate binding site |
| 573 | purification tag pos. 2 | 18 | purification tag to BiP substrate binding site |
| 573 | purification tag pos. 2 | 18 | purification tag to BiP substrate binding site |
| 447 | purification tag pos. 16 | 33 | purification tag to outside edge of BiP SBD domain beta |
| 516 | purification tag pos. 16 | 17 | purification tag to outside edge of BiP SBD domain beta |
| 579 | purification tag pos. 16 | 16 | purification tag to BiP substrate binding site |
| 581 | purification tag pos. 16 | 20 | purification tag to BiP substrate binding site |
| 581 | purification tag pos. 16 | 15 | purification tag to BiP substrate binding site |
| 585 | purification tag pos. 16 | 27 | purification tag to BiP substrate binding site |
| 620 | purification tag pos. 16 | 30 | purification tag to outside edge of BiP SBD alpha helical lid |
| 621 | purification tag pos. 16 | 25 | purification tag to outside edge of BiP SBD alpha helical lid |
| 585 | 72 | 16 | GRP94 NTD to BiP substrate binding site |
| 523 | 87 | 17 | GRP94 NTD to outside edge of BiP SBD domain beta |
| 523 | 87 | 17 | GRP94 NTD to outside edge of BiP SBD domain beta |
| 164 | 114 | 27 | interface II |
| 164 | 114 | 27 | interface II |
| 185 | 114 | 27 | interface II |
| 185 | 114 | 24 | interface II |
| 185 | 114 | 18 | interface II |
| 523 | 114 | 21 | GRP94 NTD to outside edge of BiP SBD domain beta |
| 523 | 114 | 23 | GRP94 NTD to outside edge of BiP SBD domain beta |
| 516 | 137 | 19 | GRP94 NTD to outside edge of BiP SBD domain beta |
| 553 | 137 | 16 | GRP94 NTD to outside edge of BiP SBD alpha helical domain |
| 164 | 161 | 26 | interface II |
| 164 | 161 | 21 | interface II |
| 447 | 168 | 19 | GRP94 NTD to outside edge of BiP SBD domain beta |
| 523 | 168 | 18 | GRP94 NTD to outside edge of BiP SBD domain beta |
| 164 | 169 | 15 | interface II |
| 213 | 455 | 27 | interface I |

| | | | |
|---|---|---|---|
| 213 | 455 | 24 | interface I |
| 213 | 458 | 21 | interface I |
| 446 | 458 | 18 | matches SBD orientation in cytosolic loading complex |
| 523 | 458 | 17 | matches SBD orientation in cytosolic loading complex |
| 651 | 458 | 22 | unstructured C-terminal extension of BiP to GRP94 MD |
| 352 | 473 | 22 | interface I |
| 353 | 473 | 20 | interface I |
| 352 | 474 | 17 | interface I |

**Inter-protein crosslinks of the BiP-GRP94Δ72 sample**

| BiP residue Uniprot ID P20029 | GRP94 residue Uniprot ID P08113 | Score | Crosslink outlines the following interaction |
|---|---|---|---|
| 447 | purification tag pos. 2 | 21 | purification tag to outside edge of BiP SBD domain beta |
| 585 | purification tag pos. 2 | 15 | purification tag to BiP substrate binding site |
| 620 | purification tag pos. 2 | 21 | purification tag to outside edge of BiP SBD alpha helical lid |
| 643 | purification tag pos. 2 | 23 | purification tag to outside edge of BiP SBD alpha helical lid |
| 523 | purification tag pos. 13 | 17 | purification tag to outside edge of BiP SBD domain beta |
| 446 | purification tag pos. 16 | 17 | purification tag to outside edge of BiP SBD domain beta |
| 446 | purification tag pos. 16 | 18 | purification tag to outside edge of BiP SBD domain beta |
| 447 | purification tag pos. 16 | 35 | purification tag to outside edge of BiP SBD domain beta |
| 581 | purification tag pos. 16 | 18 | purification tag to BiP substrate binding site |
| 585 | 72 | 18 | GRP94 NTD to BiP substrate binding site |
| 446 | 75 | 17 | GRP94 NTD to outside edge of BiP SBD domain beta |
| 164 | 114 | 23 | interface II |
| 185 | 119 | 15 | interface II |
| 344 | 119 | 16 | unspecific Xlink |
| 516 | 137 | 21 | GRP94 NTD to outside edge of BiP SBD domain beta |
| 521 | 137 | 17 | GRP94 NTD to outside edge of BiP SBD domain beta |
| 523 | 137 | 15 | GRP94 NTD to outside edge of BiP SBD domain beta |
| 523 | 137 | 19 | GRP94 NTD to outside edge of BiP SBD domain beta |
| 523 | 137 | 23 | GRP94 NTD to outside edge of BiP SBD domain beta |
| 213 | 161 | 24 | not compatible with any tested conformation |
| 447 | 161 | 29 | GRP94 NTD to outside edge of BiP SBD domain beta |
| 447 | 161 | 24 | GRP94 NTD to outside edge of BiP SBD domain beta |
| 523 | 161 | 21 | GRP94 NTD to outside edge of BiP SBD domain beta |
| 523 | 161 | 24 | GRP94 NTD to outside edge of BiP SBD domain beta |
| 547 | 161 | 18 | GRP94 NTD to outside edge of BiP SBD alpha helical lid |
| 164 | 168 | 25 | interface II |
| 185 | 168 | 18 | interface II |

| | | | |
|---|---|---|---|
| 523 | 168 | 25 | GRP94 NTD to outside edge of BiP SBD domain beta |
| 213 | 455 | 20 | interface I |
| 213 | 458 | 22 | interface I |
| 213 | 458 | 25 | interface I |
| 352 | 458 | 18 | interface I |
| 446 | 458 | 19 | matches SBD orientation in cytosolic loading complex |
| 447 | 458 | 20 | matches SBD orientation in cytosolic loading complex |
| 447 | 458 | 19 | matches SBD orientation in cytosolic loading complex |
| 523 | 458 | 16 | matches SBD orientation in cytosolic loading complex |
| 651 | 467 | 21 | unstructured C-terminal extension of BiP to GRP94 MD |
| 352 | 473 | 21 | interface I |
| 352 | 473 | 19 | interface I |
| 352 | 473 | 19 | interface I |
| 352 | 473 | 19 | interface I |
| 352 | 473 | 19 | interface I |
| 352 | 474 | 19 | interface I |
| 352 | 474 | 19 | interface I |
| 352 | 508 | 17 | interface I |
| 352 | 508 | 18 | interface I |
| 352 | 508 | 20 | interface I |
| 353 | 508 | 19 | interface I |
| 651 | 508 | 16 | unstructured C-terminal extension of BiP to GRP94 MD |
